# Mapping Protein Dynamics at High Spatial Resolution with Temperature-Jump X-ray Crystallography

**DOI:** 10.1101/2022.06.10.495662

**Authors:** Alexander M. Wolff, Eriko Nango, Iris D. Young, Aaron S. Brewster, Minoru Kubo, Takashi Nomura, Michihiro Sugahara, Shigeki Owada, Benjamin A. Barad, Kazutaka Ito, Asmit Bhowmick, Sergio Carbajo, Tomoya Hino, James M. Holton, Dohyun Im, Lee J. O’Riordan, Tomoyuki Tanaka, Rie Tanaka, Raymond G. Sierra, Fumiaki Yumoto, Kensuke Tono, So Iwata, Nicholas K. Sauter, James S. Fraser, Michael C. Thompson

## Abstract

Understanding and controlling protein motion at atomic resolution is a hallmark challenge for structural biologists and protein engineers because conformational dynamics are essential for complex functions such as enzyme catalysis and allosteric regulation. Time-resolved crystallography offers a window into protein motions, yet without a universal perturbation to initiate conformational changes the method has been limited in scope. Here we couple a solvent-based temperature jump with time-resolved crystallography to visualize structural motions in lysozyme, a dynamic enzyme. We observed widespread atomic vibrations on the nanosecond timescale, which evolve on the sub-millisecond timescale into localized structural fluctuations that are coupled to the active site. An orthogonal perturbation to the enzyme, inhibitor binding, altered these dynamics by blocking key motions that allow energy to dissipate from vibrations into functional movements linked to the catalytic cycle. Because temperature-jump is a universal method for perturbing molecular motion, the method demonstrated here is broadly applicable for studying protein dynamics.

## Main

Conformational dynamics drive ubiquitous protein functions, such as enzyme catalysis, signal transduction, and allosteric regulation^1^. The study of protein motions is critical to our mechanistic understanding of these fundamental biological phenomena, and the ability to rationally control protein motions is a frontier for protein engineers who seek to design biomolecules with increasingly complex functions. Nevertheless, accurately modeling protein motions with atomic resolution remains a longstanding challenge for the field of structural biology^2^. This challenge arises because data from imaging techniques such as X-ray crystallography and cryoEM are inherently ensemble-averaged over space and time^3^. Thus, information about spatiotemporal coupling between observed alternate conformations is lost, and intermediate states that are only transiently populated at equilibrium remain invisible. For this reason, high-resolution structural information is often combined with spectroscopic measurements. Despite the power of such integrative approaches, it can be difficult to correlate spectroscopic observables with specific features of molecular structures. Time-resolved X-ray crystallography (TRX) overcomes these challenges by obtaining high-resolution information in both the spatial and temporal domains^4–6^. In this pump-probe technique, a rapid perturbation drives molecules out of conformational equilibrium, then structural snapshots are captured as the ensemble relaxes to a new equilibrium. By sampling a series of pump-probe time delays, kinetic couplings between conformers and transiently-populated structural states can be observed.

TRX experiments were pioneered on photoactive proteins^6–8^ and have become more readily accessible with the advent of X-ray free-electron lasers (XFELs), brighter synchrotrons, and serial crystallography^9–11^. The scope of TRX has broadened further still through the use of photocaged ligands^12^, rapid mix-and-inject experiments^13^, and electric-field-based perturbations^5^. Still, it remains challenging to excite a protein’s intrinsic dynamics in a generalizable manner for time-resolved structural studies. While many perturbations are tailored to the protein of interest, protein dynamics are universally coupled to thermal fluctuations of the surrounding solvent^14^. Recent multi-temperature-crystallography experiments demonstrated that modifying temperature allows one to tune the conformational ensemble sampled at equilibrium, paving the way for temperature-jump TRX experiments^15, 16^. Infrared laser-induced temperature jumps (T-jump) are routinely used to study protein folding and enzyme dynamics^17–19^. In such experiments, a mid-IR laser excites the O-H stretching mode of water molecules, resulting in rapid heating of the sample. Both solvent heating^20^ and heat transfer through the protein^21^ happen on faster timescales than most functional protein motions, removing the conformational ensemble from thermal equilibrium. Subsequently, the system relaxes to a new equilibrium, determined by the final temperature, as a subset of the molecules in the sample populate higher-energy conformational states. At both temperatures, the protein exists as an ensemble of conformations. At higher temperatures, higher energy conformations will be more significantly populated due to Boltzmann statistics^3, 22^. Previously, T-jump was coupled to small-angle X-ray scattering (SAXS) to perform time-resolved measurements of structural dynamics^23, 24^. Infrared laser pulses have also been used to study thermal denaturation of an enzyme *in crystallo,* but prior work lacked rigorous time-resolved measurements of the motions induced by rapid heating^25^.

Here, we present time-resolved temperature-jump serial femtosecond crystallography (SFX) experiments on a model system, lysozyme, demonstrating that T-jumps are an effective perturbation for measuring the intrinsic conformational dynamics of proteins. We used a nanosecond pulsed laser tuned to the mid-IR region of the electromagnetic spectrum to heat the bulk solvent within and around lysozyme microcrystals. These microcrystals were delivered to the pump-probe interaction region using a free-flowing microfluidic stream and were subsequently probed using ultrafast, high-intensity X-ray free-electron laser (XFEL) pulses (**Figure 1**). We detected the introduction of a T-jump directly from singular value decomposition (SVD) of the isotropic diffuse scattering present in diffraction images, and using a repertoire of real space analysis tools including weighted difference electron density maps^26^ and structure refinement against extrapolated structure factors^27, 28^, we revealed that the rapid heating of the crystals excites the intrinsic dynamics of crystallized lysozyme molecules.

**Figure 1.**
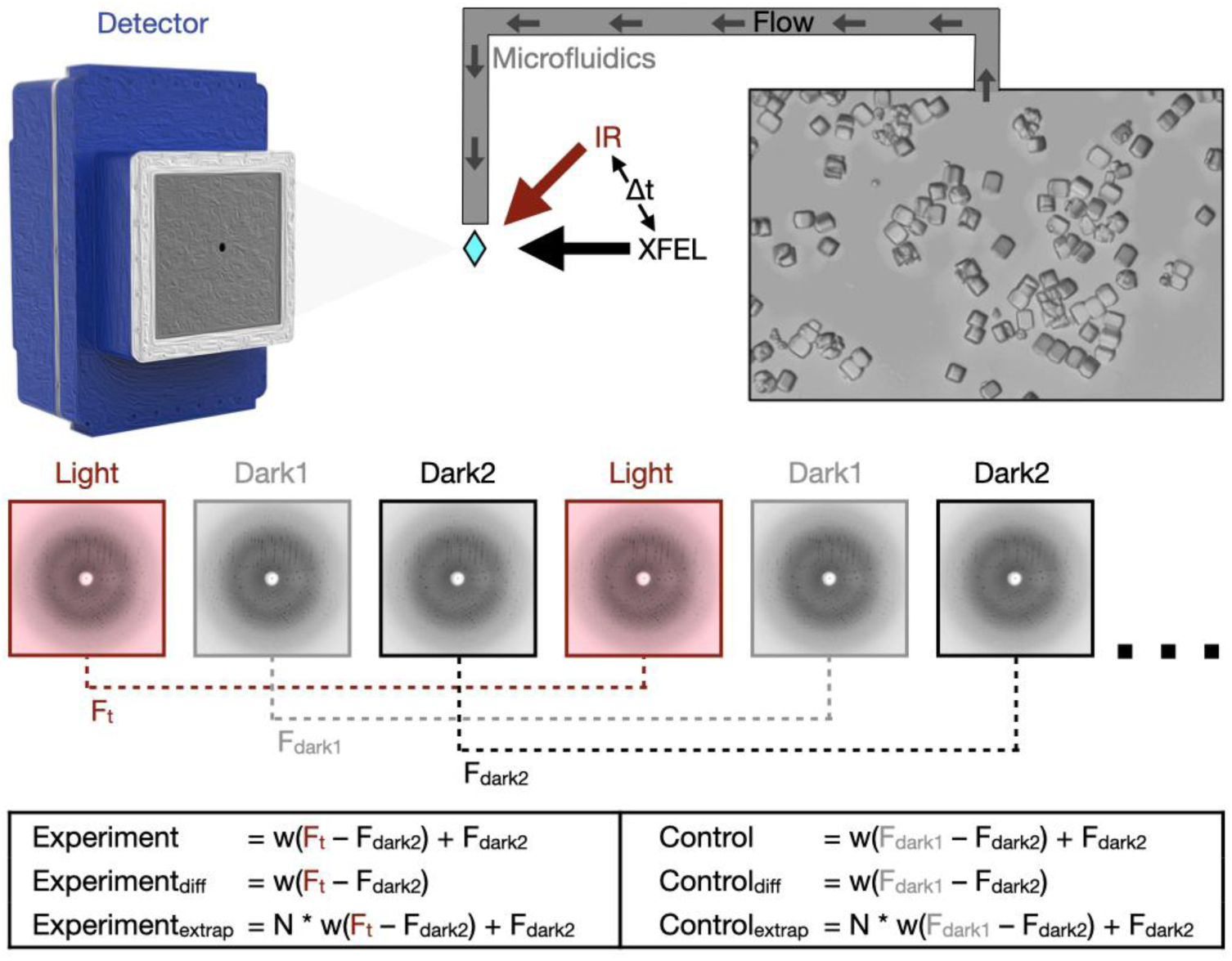
Schematic of T-jump TRX experiment. Lysozyme crystals were delivered to the pump-probe interaction region via a microfluidic jet. Light and dark images were collected in an interleaved manner, with light images defined as those where the crystal was pumped with an IR laser at a defined time-delay (Δt) prior to being probed by the XFEL. Dark images were collected with the IR pump shutter closed and an XFEL probe. These images were combined in post-processing to create a set of structure factors for each time delay (F_t_) and as well as two corresponding sets of dark structure factors (F_dark1_ and F_dark2_). Experiments and matching controls as defined above and in the text were analyzed to identify time-dependent structural changes.

At short pump-probe time delays (20 ns), we observe the signatures of widespread atomic vibrations, which dissipate to reveal coordinated motions in functionally-relevant regions of the enzyme structure at longer pump-probe time delays (20-200 μs). Our observations are consistent with prior work to elucidate the role of conformational dynamics in lysozyme’s function. Furthermore, T-jump TRX allowed us to directly visualize the effect of an inhibitor on the observed motions of the enzyme. This work opens the door for future T-jump TRX experiments on diverse protein systems by leveraging the inextricable connection between temperature and macromolecular dynamics. This includes proteins whose activities cannot be triggered by light or rapid mixing.

### Instrumentation and Data Collection for T-Jump Crystallography

We paired an IR laser with high-resolution serial femtosecond crystallography (SFX)^29^ in a pump-probe configuration to measure the conformational dynamics of lysozyme in real-time and at atomic resolution. Leveraging instrumentation compatible with standard photoactive TRX experiments^30, 31^; we tuned the pump laser to the mid-IR range to excite the O-H stretch mode of water with a pulse duration of approximately 7 ns, resulting in a T-jump^32^. For SFX measurements, we embedded batch-grown lysozyme microcrystals in an 18% hydroxyethylcellulose carrier medium^33^ (**Table EX1)** and delivered them to the pump-probe interaction region using a viscous extrusion device^34^ that formed a free-flowing microfluidic jet (**Figure 1**). Once delivered to the interaction region, crystals were rapidly heated by the IR laser. At defined time delays following the IR heating pulse (20 ns, 20 μs, and 200 μs), we probed the sample with ultrafast, high-brilliance X-ray pulses (**Table EX1**) from the SPring-8 Angstrom Compact Linear Accelerator (SACLA) XFEL^35^, and measured X-ray diffraction using a custom multi-panel CCD detector^36^. The fastest measurable time delay was 7 ns, limited by the duration of the mid-IR pump laser pulse. In each of our datasets, T-jump measurements were interleaved with two measurements in which the IR laser shutter was closed, providing internal controls (**Figure 1**). We refer to the sequential interleaved dark measurements as “dark1” and “dark2”, and these sets of interleaved control measurements were treated as separate datasets. Therefore, each of our T-jump datasets (20 ns, 20 us, 200 us) is paired with two internal control datasets (F_dark1_ and F_dark2_). Additionally, a dataset was collected where the IR laser shutter remained closed (no interleaved T-jumps) which we refer to as F_laser off_.

### T-Jump Detection from Diffuse Scatter

During the experiment, we performed online data analysis with a computational pipeline developed specifically for data collection at SACLA^37^, which identified, in real-time, detector images that contained X-ray diffraction from crystals and subsequently integrated and merged the growing dataset to provide preliminary feedback on data quality and dataset completeness. Following the experiment, data were processed with more rigorous optimization of parameters used for indexing and integration, as well as post-refinement during merging^38–41^. A summary of the data quality is available (**Tables EX1, EX2 & EX3**), and a detailed description of both online and offline data processing procedures are provided (**Methods**).

To confirm that the application of the IR pump laser introduced a temperature jump in the sample, we modified tools from previous SAXS experiments^23^ to analyze the scattering signal resulting from the solvent around and within the protein crystals. Starting from raw diffraction images identified as crystal hits, we azimuthally-averaged the total scattering and applied a median filter to remove the effect from Bragg peaks. This generated a series of one-dimensional scattering curves which contained the isotropic signature from bulk water. These scattering curves were scaled and combined in a matrix that was analyzed using singular value decomposition (SVD). SVD generated a set of right singular vectors (basis vectors) (**Figure 2a**) and a set of left singular vectors that quantified the contribution of each basis vector to every scattering curve used to construct the input matrix (**Figure 2b**). We observed that the contribution of the right singular vector associated with the second largest eigenvalue (v1) to each scattering image was highly correlated with whether the pump laser pulse had been applied, as determined by the distribution of entries in the corresponding left singular vector (**Figure 2c**). In contrast, the contribution of other basis vectors identified by SVD (v0, v2, v3 in Figure 2) do not show the same dependence on the laser status. This observation provided a means to retrospectively confirm the introduction of a T-jump in all images collected under pump laser illumination by directly analyzing laser-dependent changes in the background solvent scattering signal.

**Figure 2.**
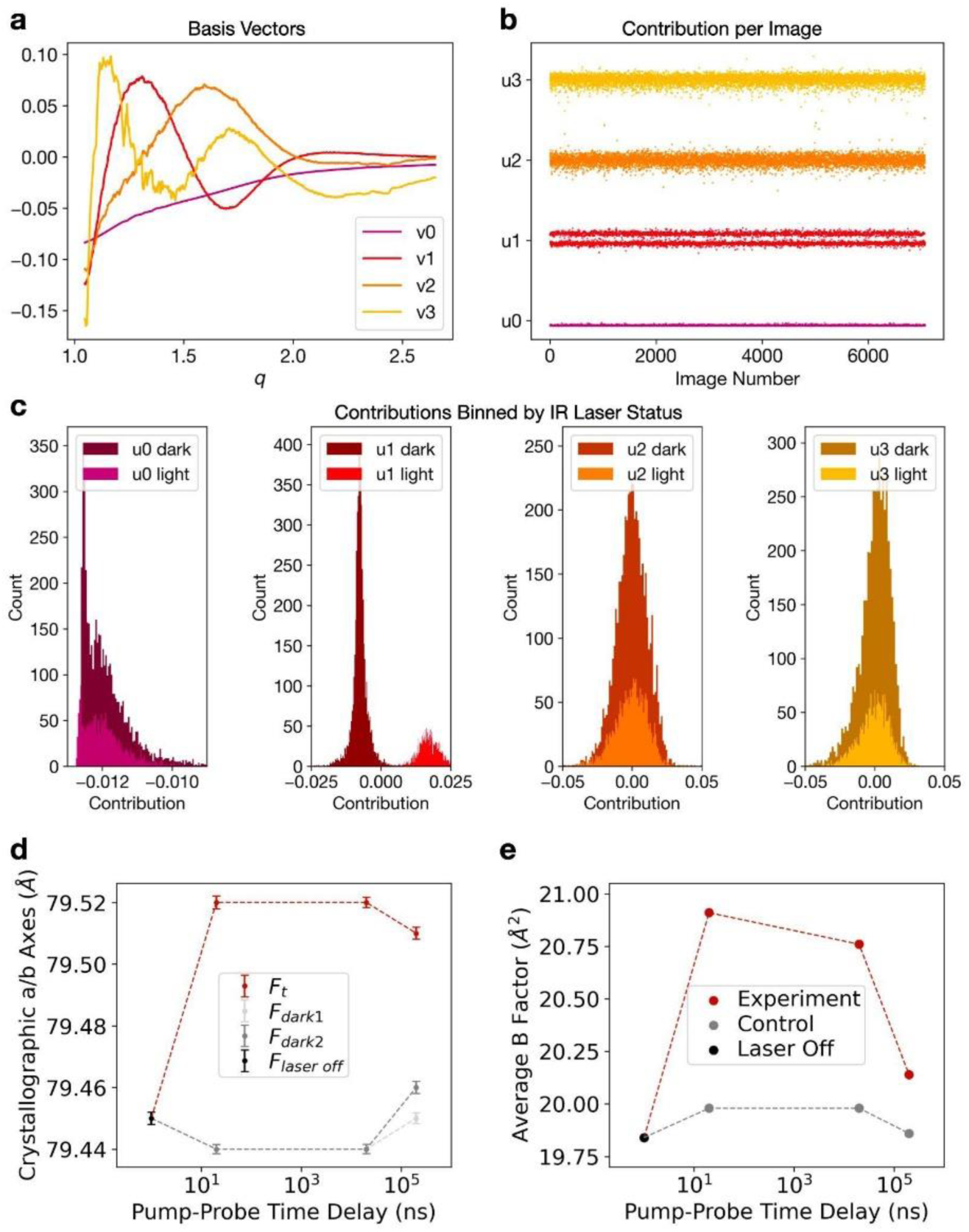
Experimental detection of temperature-jump. Radial averages of diffraction images from the 20ns dataset were decomposed using SVD. (**a**) The basis vectors (vn) associated with the largest four singular values were visualized, as was the contribution (un) of each corresponding basis vector to each radial average (**b**). U1 values were split, which was clarified by plotting the values as histograms (**c**). The values and sign of u1n associated with the second largest singular value, correlate with IR laser status, as reported by the IR laser diode. (**d**) Analysis of the crystallographic a/b axis lengths, equivalent under P43212 symmetry, as a function of pump-probe time delay reveal thermal expansion of the unit cell. All values in (**d**) are mean ± 95%CI. Time zero corresponds to the laser-off (laser off) dataset, with laser-on (light) and interleaved laser-off (dark1 & dark2) datasets plotted for each pump-probe time delay. Unit cell dimensions expand following perturbation with an IR laser, while unilluminated data show consistent unit cell dimensions over the course of the experiment. (**e**) Similarly, the average B-factor of refined models increases following perturbation with an IR laser. Models were refined against Laser Off, Experiment, or Control structure factors.

### T-Jump Induces Unit Cell Expansion and Increases Global Protein Dynamics

Using the processed diffraction data, we compared the unit cell dimensions as a function of pump-probe time delay. We observed that application of the IR laser results in rapid expansion of the unit cell (**Figure 2d**). This thermal expansion corresponds to <1% of the length of the unit cell axes yet is statistically significant because each serial crystallography dataset contains >10,000 unique crystal measurements (**Tables EX2, EX4**). Furthermore, we observed that the average unit cell parameters for the interleaved F_dark1_ and F_dark2_ datasets matched the F_laser off_ dataset much more closely than the unit cells seen in F_t_ datasets. This comparison suggests that the observed unit cell expansion is due to the application of the pump laser rather than variations in other parameters, such as crystal batch or sample delivery conditions. The thermal expansion of the unit cell provides further evidence of a laser-induced T-jump.

As an initial step toward identifying structural changes that were induced by a T-jump, we refined models against the F_t_ structure factors from each time delay. The resulting atomic coordinates were nearly identical to one another as well as to a structure refined against the F_laser off_ data; however, we did observe an increase in the average atomic B-factor in the T-jump structures (**Figure 2e**). Collectively, these observations indicate small populations of high-energy conformational states and/or motions that are well-modeled by the harmonic approximation of the B-factor, especially for short (20 ns) pump-probe time delays. The models refined against the raw F_t_ structure factors were deposited to the Protein Data Bank (PDB).

### Time-Resolved Electron Density Changes

To gain further insight into time-resolved changes in the protein structure, we created a series of difference electron density maps. These maps were created by subtracting interleaved dark2 measurements (F_dark2_) from T-jump measurements (F_t_) combined with phases from the corresponding dark state models as refined against (F_dark2_) data. Additionally, we applied a weighting scheme developed to improve the estimation of difference structure factors calculated from noisy data^26^. We viewed the resulting maps in the context of the corresponding dark state model and observed features consistent with time-dependent changes in the enzyme’s conformational ensemble (**Figure 3a**). Generally, we observed that time-resolved difference electron density evolved from a strong ubiquitous signal centered on atom positions to coordinated positive and negative peaks that suggest correlated motions within the enzyme. Features appearing at the 20 ns time delay were widespread, with negative peaks centered on heavy atoms (non-hydrogens), surrounded by distinct positive halos (**Figure 3a, Figure EX1a, Video S1**). In difference electron density maps corresponding to microsecond time delays (20 and 200 us), most difference density features manifested as coupled positive and negative peaks adjacent to the protein molecule. At 20 µs, the largest peaks appeared at sparse locations adjacent to the backbone, indicating the beginning of coordinated motions (**Figure 3a, Video S2**). These motions continued to evolve into paired positive and negative peaks indicating coordinated shifts of the backbone away from the conformation that is most prevalent in the dark state at the 200 µs pump-probe time delay. Residues 97-100 provide a clear example of such motions (**Figure 3a, Video S3**).

**Figure 3.**
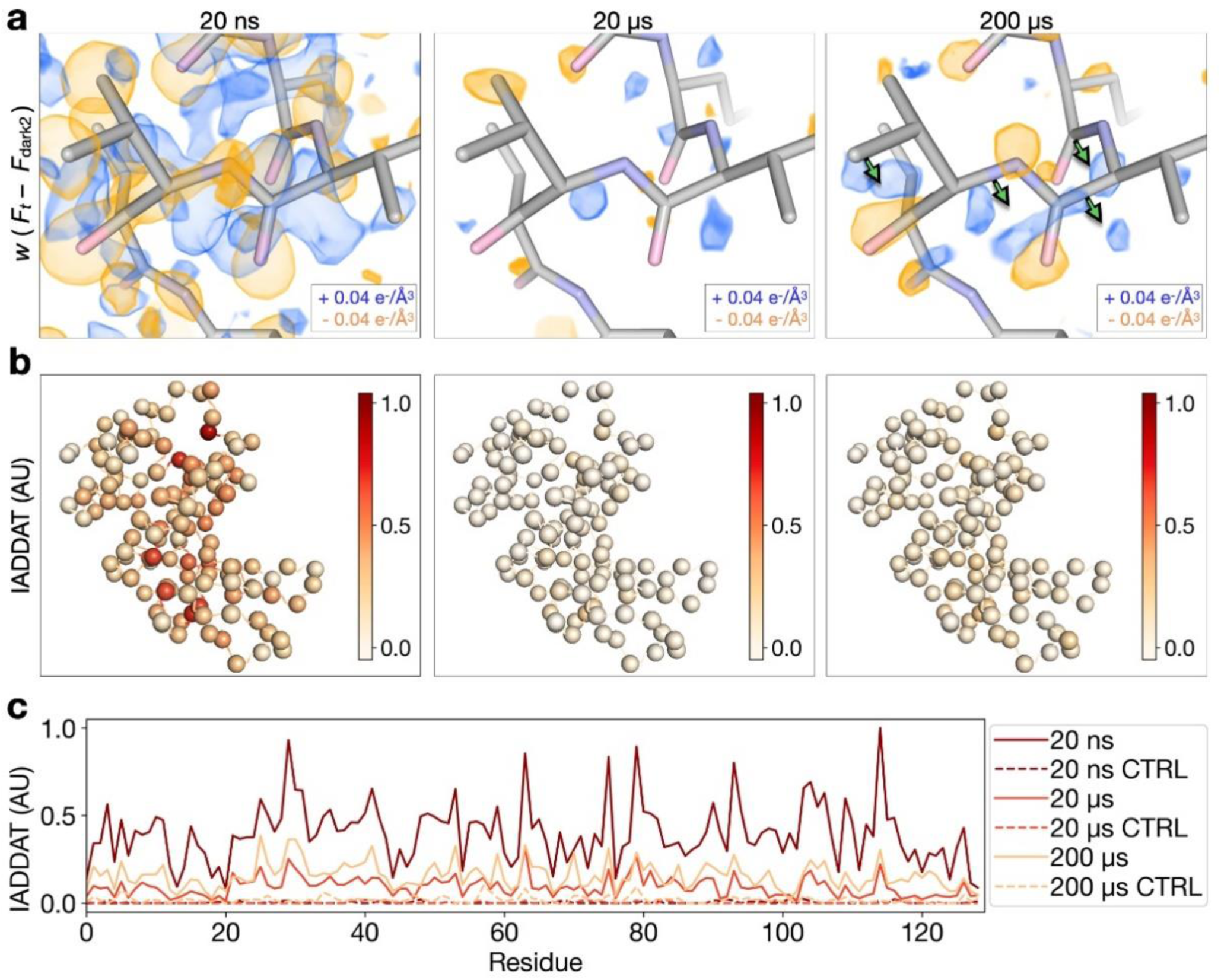
Time-resolved difference electron density evolves over time following T-jump. (**a**) Comparison of weighted difference electron density maps (Ft – Fdark2) for each pump-probe time delay, centered around residues 97-100. Maps were visualized at an absolute contour level of ± 0.04 e^-^/Å^3^ alongside corresponding refined models. While model coordinates appeared stable across pump-probe time delays, difference maps revealed time-resolved changes to the T-jump induced signal, with evidence for coordinated motions (green arrows) apparent by 200 µs. (**b**) IADDAT was calculated as an average value per residue for each pump-probe time delay, then mapped onto C-alpha positions (spheres) of the respective model, and plotted as a function of residue number (**c**). Comparison of IADDAT values for experimental maps relative to matched controls revealed low levels of noise across the series.

To test whether the peaks observed in our difference maps were driven by the T-jump rather than experimental noise, we calculated a series of matching control difference maps using the interleaved dark datasets (F_dark1_ and F_dark2_) (**Figure 1**). Control maps contained very few features, confirming that the signals observed in the time-resolved maps were far above the threshold of experimental noise (**Figure EX1a**). To quantify this comparison, we calculated pairwise real space correlation coefficients (RSCC) between all difference maps (**Figure EX1b**). We expected maps dominated by random noise to display no correlation, whereas systematic variations or common signals in the map would drive the RSCC towards 1. Pairwise comparisons between control maps revealed RSCC values near 0, indicating the presence of noise alone. Control maps compared to experimental maps revealed RSCC values again near 0, with a minor increase in matched pairs, indicating minor experimental variations common to the interleaved data. Finally, pairwise comparisons between experimental maps showed much larger RSCC values, ranging between 0.3 and 0.7, indicating common signals distributed over time.

To further characterize the spatial distribution of features observed in our time-resolved difference maps we integrated the absolute difference density above a noise threshold (IADDAT)^5, 42^. In this calculation, difference peaks with an absolute value greater than 0.04 e^-^/Å^3^ and within 2.5 Å of an atom (excluding waters) are summed, and an average value is calculated on a per residue basis. Then, the per-residue values are scaled and mapped onto corresponding C-alpha positions (**Figure 3b**). At all pump-probe time delays, the distribution of difference density is non-uniform across the molecule, indicating specific regions of enhanced dynamics (**Figure 3b**). The difference density is also quantitatively strongest in the map corresponding to the 20 ns pump-probe time delay and weakest in the map corresponding to the 20 us time delay (**Figure 3c**). This spatial analysis revealed which local regions of the enzyme respond most strongly to the T-jump, as well as the timescales of these responses. IADDAT calculations on matched control maps (F_dark1_ -F_dark2_), revealed virtually no signal.

### Modeling Time-Dependent Conformational Changes

Given the time-resolved changes evident in difference electron density maps, we next sought to model the specific structural changes that give rise to those signals. Because the nature of difference density signals differed substantially as a function of pump-probe time delay, we used distinct approaches to model signals appearing on the nanosecond and microsecond timescales.

As noted, the dominant difference density features that we observed at the 20 ns pump-probe time delay manifested as negative peaks over atomic positions, surrounded by positive halos (**Figure 3a, Figure EX1a, Video S4**). We hypothesized that these features corresponded to an overall increase in atomic B-factors resulting from increased thermal motion following T-jump, because the subtraction of a taller but narrower gaussian function from one that has the same area under the curve but is shorter and broader would produce a difference function with a similar shape. To test this hypothesis, we used two sets of structure factors calculated from our dark state model. Specifically, we calculated one set of structure factors directly from the atomic coordinates and refined B-factors, and a second set of structure factors from the same coordinates, but with B-factors that had been scaled by a factor of 1.2, representing a 20% increase. We then used these sets of calculated structure factors to generate a difference map showing how an increase in B-factors due to a T-jump would manifest in the electron density, and noted a striking similarity with the experimentally-derived map (**Figure EX2, Video S5**). We determined the real space correlation coefficient between this hypothetical map and the map calculated from our 20 ns T-jump data and found extremely good agreement (CC=0.67, **Figure EX1b**). Systematic comparison of this simulated difference map to experimentally-derived difference maps, via real space correlation coefficients, revealed additional similarities (**Figure EX1b**). The simulated map has a weaker, but still significant, correlation with 20 us and 200 us experimental difference maps (0.33 and 0.37, respectively), likely corresponding to negative peaks appearing at atom positions as electron density moves away from these locations.

Next, using difference maps as a guide, set at a contour level of 0.04 e^-^/Å^3^ to match our IADDAT analysis, we manually built alternative conformations for several regions of the protein where the difference density could be interpreted explicitly, including residue 23 (inspired by the 20 ns difference map) as well as a short loop composed of residues 97-100 which lies at the end of an α-helix adjacent to the active site (inspired by the 200 µs difference map) (**Figure 4a**). We included these alternative conformations in a multi-conformer model, and conducted an additional series of refinements against “extrapolated” structure factor magnitudes (ESFMs)^27, 28^ to assess whether the alternative conformations could represent high-energy states of the enzyme. The procedure used to generate datasets containing ESFMs is described in Figure 1 and Equation 2, and has been implemented by others^5, 6^. Briefly, we multiplied the weighted experimental structure factor difference, w(F_t_ – F_dark2_), for a given pump-probe time delay by an arbitrary extrapolation factor (N), and then added that product to the structure factors measured for the dark state (F_dark2_). This manipulation of the experimental data effectively scales up the contribution of high-energy states of the molecule that are populated by T-jump to the overall structure factor magnitudes. For each pump-probe time delay in our T-jump series, we refined our model, containing the dark state structure and potential high-energy conformations, against a series of ESFM sets generated with increasing extrapolation factors (**Methods**). Following refinement of coordinates, B-factors, and occupancies, we observed that the major dark state conformation was unperturbed and that the newly built conformations moved very little (**Figure 4a, Videos S6 and S7**). We investigated the occupancies of these conformations as a function of extrapolation factor (**Figure 4b**), and noticed that occupancies of the novel conformers generally increased as a function of extrapolation factor. This observation further confirms the correlation between the modeled molecular motions and laser-induced changes to the structure factors, but also suggested that only a very small fraction of protein molecules have transitioned to higher energy conformational states, because we were not able to extrapolate enough to model the high-energy state at full occupancy. The need to apply large extrapolation factors, combined with the fact that temperature-induced conformational changes are dispersed throughout the unit cell rather than concentrated near a chromophore, limit these refinements against ESFMs^27^.Nevertheless, using this approach, we could detect specific molecular motions, including short amplitude motions (e.g. rotation of Tyr23) that were most evident at the 20ns pump-probe time delay, and larger motions (e.g. the backbone shift of loop 97-100) that were most evident in the 200 µs dataset. The backbone shift that we detected for residues 97-100 in the 200 µs difference electron density map was consistent with backbone motion calculated by normal mode analysis of the ground state structure (**Figures 4a and EX6**).

**Figure 4.**
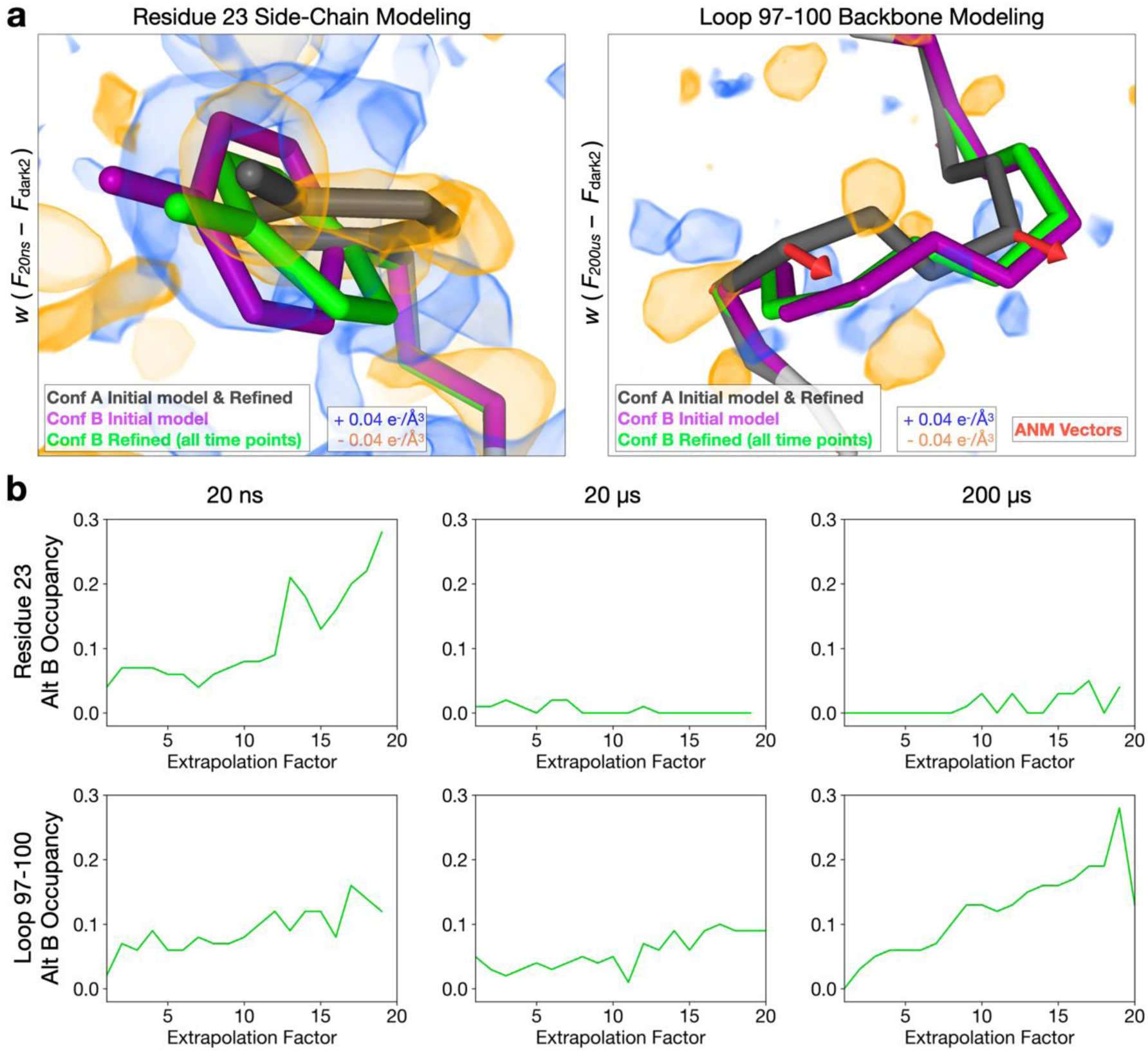
Explicit modeling of time-resolved structural changes. (**a**) For all pump-probe time delays, difference electron density maps were visualized at an absolute contour level of ± 0.04 e^-^/Å^3^ alongside the refined models. Alternate conformations were manually modeled into the experimental difference density for several regions, including residues 23 and 97-100, then refined against extrapolated structure factor magnitudes. An anisotropic network model (ANM) was developed based on the ground state (conformation A of the initial model) and vectors were visualized for comparison with alternate conformers. (**b**) Occupancies of the new conformations were examined as a function of increasing extrapolation factors. The stability of the hypothetical high-energy states during the coordinate refinement, and the increase in their occupancies with increasing extrapolation factor provide evidence that these conformations are populated in the ensemble.

### Inhibitor Binding Alters T-Jump Induced Dynamics

In addition to the datasets described above, we also examined analogous data for lysozyme bound to a naturally-occurring inhibitor, chitobiose, which is known to bind to the active site and stabilize a “closed” form of the enzyme (**Figure 5a**). Specifically, for chitobiose-bound crystals, we collected T-jump data corresponding to two pump-probe time delays (20 ns and 200 μs, see **Table EX4**). Time-resolved difference maps calculated for chitobiose bound crystals revealed an altered dynamic response to T-jump relative to the apo crystals (**Figure 5b**). Qualitatively, while the 20 ns difference density looks similar for both apo and chitobiose-bound crystals, marked differences are evident by 200 µs. In the case of the apo enzyme, features in the time-resolved difference maps change substantially between 20 ns and 200 µs. In contrast, when chitobiose is bound, the features in the time-resolved difference maps change relatively little between 20 ns and 200 µs (Figure 5b). These observations were confirmed quantitatively using IADDAT calculations. Specifically, we looked at the overall difference in IADDAT (|ΔIADDAT|) between the 20 ns and 200 µs time delays for both the apo and chitobiose-bound crystals (Figure 5c & 5d). We observed that the difference in IADDAT as a function of pump-probe time delay is much more pronounced for the apo enzyme relative to when the inhibitor is bound. RSCC calculations further confirmed these observations, showing high pairwise correlations between the apo 20 ns, chitobiose-bound 20 ns, and chitobiose-bound 200 µs difference maps, each of which have a much weaker correlation with the apo 200 µs difference map (Figure EX1b).

**Figure 5.**
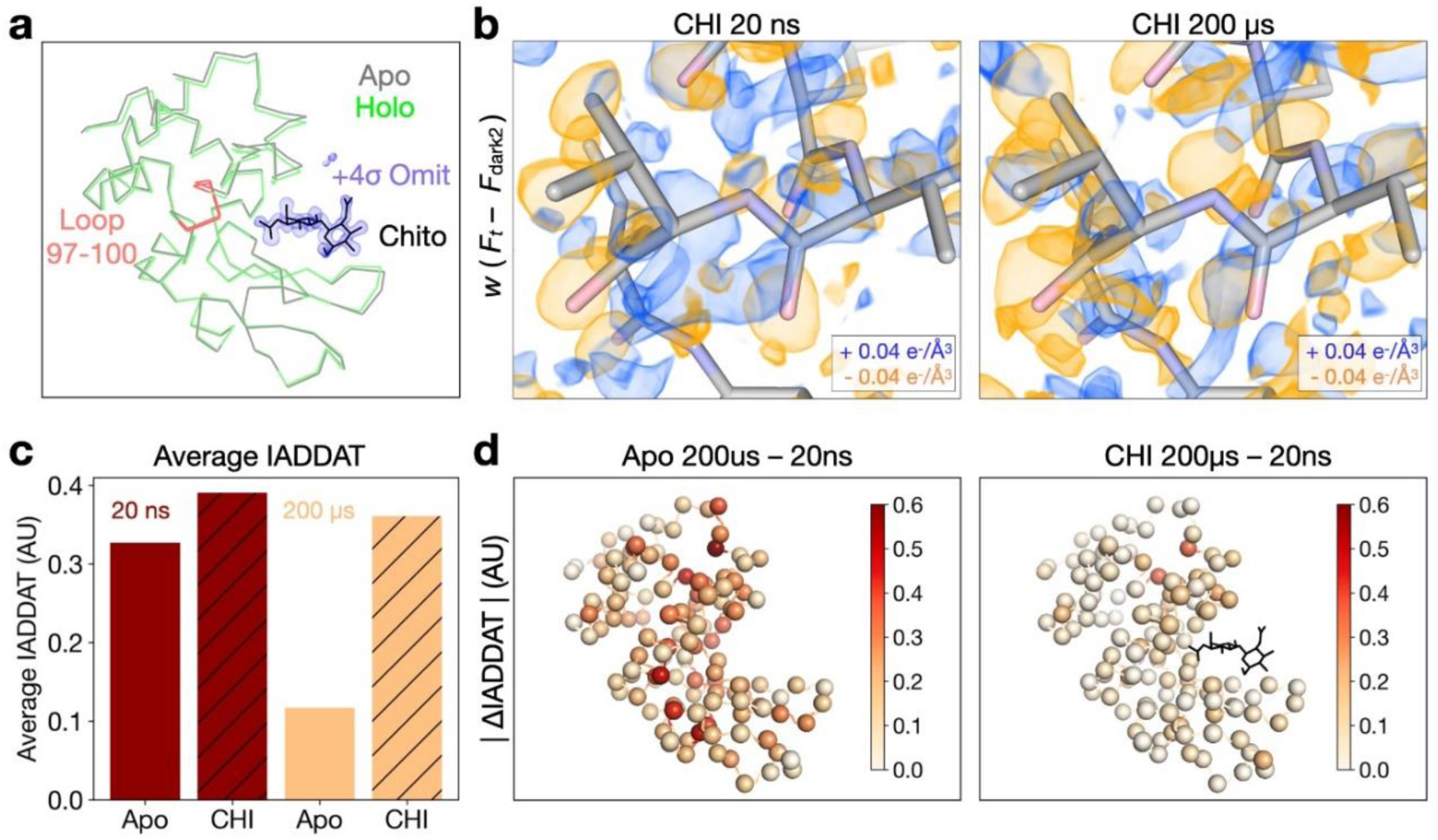
Chitobiose binding perturbs changes induced by T-jump. (**a**) Ribbon diagrams of lysozyme structures (laser off) in the apo (gray) and chitobiose-bound (holo, green) forms show a decrease in distance between the two lobes of the protein upon ligand binding, characteristic of the active site “closing” motion. Chitobiose is shown as black sticks, along with a ligand omit map contoured to +4σ and carved within 5 Å of the chitobiose molecule. (**b**) Visualization of weighted difference density maps (Ft – Fdark2) for chitobiose-bound data sets show similar in the apo and ligand-bound states. Maps were visualized at an absolute contour level of ± 0.04 e-/Å^3^ alongside initial refined models for each pump-probe time delay. (**c**) Comparing average IADDAT values for ligand-bound and apo maps revealed similar signal levels at 20 ns, with substantial differences appearing by 200 µs. (**d**) Mapping absolute differences in IADDAT between 20 ns and 200 µs (|ΔIADDAT|) values onto the structure (C-alphas as spheres) further reveals that time-resolved changes are more pronounced for the apo enzyme than for the inhibitor-bound enzyme.

## Discussion

The results presented here demonstrate that infrared laser-induced temperature-jump can be used as an effective perturbation method for time-resolved X-ray crystallography to map the dynamics of a biological macromolecule at atomic resolution. We performed pump-probe serial femtosecond crystallography using a nanosecond pulsed laser, tuned to mid-IR wavelengths (1443 nm), to rapidly heat the solvent surrounding and permeating microcrystals of the dynamic enzyme lysozyme (40.7% solvent), and monitored the resulting molecular motions using ultrafast X-ray pulses from an X-ray free-electron laser. Using algebraic tools, leveraged from previous solution X-ray scattering (SAXS/WAXS) experiments^23^, we detected the introduction of a T-jump, which was corroborated by observations of unit cell thermal expansion and increased atomic displacements (higher refined B-factors). Next, we implemented a series of analyses common in time-resolved protein crystallography, including calculation of uncertainty-weighted difference electron density maps^26^, spatial integration of difference electron density^5, 42^, and refinement against extrapolated structure factor magnitudes^27, 28^, allowing us to identify high-energy conformational states of the protein, and to estimate the timescales on which they are populated.

The goal of a T-jump crystallography experiment is to map the structural dynamics of the crystallized molecule. We selected lysozyme as our subject for this initial study, because it crystallizes easily, diffracts well, and is known to undergo a characteristic hinge-bending motion that results in domain closure around the active site^43, 44^. Although limited in temporal resolution, the data we collected and the analyses we performed provide a coarse-grained model for the evolution of conformational dynamics in lysozyme following T-jump that is consistent with published literature on lysozyme dynamics^43, 45^. Our analysis of experimental maps supports a rapid increase in atomic displacement parameters (B-factors) within 20 ns of the pump laser pulse, evidence of increased harmonic motion resulting from heat transfer to the protein from the thermally-excited solvent. The spatial distribution of difference electron density features at very short pump-probe time delays (e.g. 20 ns) is not uniform (**Figures 3a, 3b, 5b & 5d**) suggesting that heat flows more readily into some regions of the lysozyme molecule than others. We also modeled high-energy conformations directly from time-resolved difference electron density maps and refined them against extrapolated structure factor magnitudes. This analysis revealed motions such as rotamer flips on fast timescales (e.g. 20 ns), and larger motions, including correlated shifts of loops spanning multiple residues, on slower timescales (e.g. 200 µs). The most noticeable of these movements includes a short loop encompassing residues 97-100, which is at the end of an α-helix adjacent to the active site and known to be mobile during lysozyme hinge bending^46^ (**Figure 5a**). These observations yield a model wherein T-jump first excites harmonic thermal motions and short-amplitude anharmonic motions, such as rotamer flips, on the nanosecond time scale. In a small population of molecules, these fast motions are subsequently converted to larger-scale conformational changes that are linked to the catalytic cycle, such as loop displacements (e.g residues 97-100) that occur on the microsecond (or longer) timescale and are coupled to the active site (**Figure 6**)^47^. Our observations are in agreement with a variety of other kinetic measurements performed on the architecturally similar, but evolutionarily unrelated, T4 lysozyme^48^, including single-molecule techniques, that demonstrated protein motions with transition times in the range of tens to hundreds of microseconds^49–51^. Notably, the intermediate pump-probe time delay (20 µs) in our limited time series contains the least evidence for high-energy conformations in both IADDAT and ESFM refinement analyses. The lack of strong difference density at this pump-probe time delay likely results from a mixture of high-energy conformational states being populated within tens of microseconds, before the ensemble narrows once again as the system approaches a new thermal equilibrium within hundreds of microseconds. This notion is consistent with single-molecule measurements that revealed large amplitude motions in lysozyme occur along multiple distinct trajectories with different transition times and intermediates^52^.

**Figure 6.**
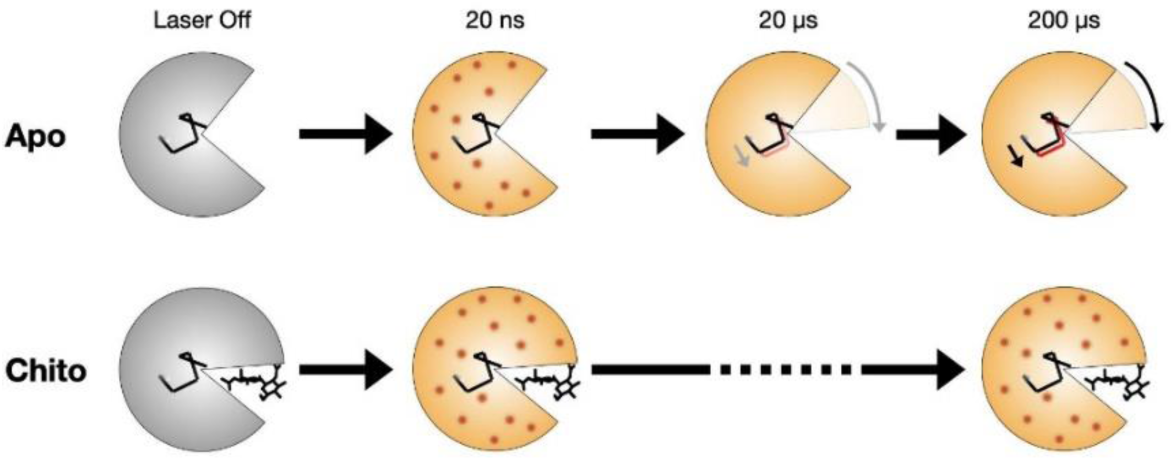
Schema of time-resolved conformational changes in lysozyme following T-jump. The cartoon highlights closure of the active site cleft upon chitobiose binding, with subsequent representations of time-resolved structural changes following T-jump. At short pump-probe time delays (20 ns) atomic vibrations (shown as red dots) are present in both the apo and inhibitor-bound structures. These vibrations persist in the inhibitor-bound structure but dissipate into more complex motions in the apo structure, including the bending of a helix that lies at the hinge point of the lysozyme molecule.

To assess whether our analysis detected functional protein motions, rather than spurious thermal fluctuations, we performed T-jump measurements in the presence of an orthogonal perturbation to the enzyme’s functional dynamics; specifically, the binding of chitobiose. Chitobiose is a natural inhibitor of lysozyme that mimics the bound substrate and shifts the enzyme’s conformational equilibrium toward the more compact “closed” state in which the two lobes of the enzyme clamp down on the carbohydrate in the active site^43, 44^. We observed, both visually and quantitatively using real-space map correlations (**Figure EX2b**), that difference electron density maps corresponding to the 20 ns pump-probe time delay look similar for both the apo and chitobiose-bound data sets, demonstrating that the fast onset of short-amplitude motions is similar in both cases. However, at the long time delay (200 µs), the difference electron density maps calculated from the chitobiose-bound data do not show evidence for dissipation of short amplitude motions into larger-scale conformational changes, as seen for the apo enzyme, but instead, remain similar to the maps corresponding to short (20 ns) pump-probe time delays (**Figure EX1b**). Additionally, the B-factors of the chitobiose-bound model remain high when refined against data corresponding to the 200 µs time delay relative to the apo model (**Figure EX3**). The inhibitor is known to restrict larger-amplitude motions (e.g. hinge bending) related to substrate binding and catalysis, locking it in a “closed” conformation. Accordingly, the presence of the inhibitor does not affect the rapid onset of thermal motion through the enzyme, but it does abolish the time-resolved changes that we attribute to microsecond functional motions (**Figure 6**), likely pushing them to the millisecond or longer timescale, outside the window of observation for our experiment. Corroborating our model, NMR spectroscopy has shown that binding of a similar carbohydrate inhibitor (chitotriose) to lysozyme has little effect on the global ps-ns motions^47^, while FRET experiments reveal that inter-domain dynamics are slowed substantially in the presence of substrate.

The work described here demonstrates that T-jump can be used as an effective perturbation to resolve conformational dynamics at atomic resolution using TRX. While limited in temporal resolution, our results allow the coarse mapping of dynamics in hen egg-white lysozyme (HEWL) across four logarithmic decades of time. While the structure of HEWL has been studied extensively by X-ray crystallography for many years, most experiments that have measured lysozyme dynamics with kinetic detail have been performed on the analogous T4 lysozyme^48^. Our observations begin to close this gap by revealing dynamics on similar timescales in the two enzymes. Because temperature change is a universal method to perturb molecular motion, we anticipate that T-jump crystallography will be applied to other systems that can undergo functional conformational changes in the crystal lattice, opening the door for broad exploration of macromolecular dynamics. While our initial experiments offer promise for the use of T-jump as a perturbation in TRX, they also highlight some of the key challenges that remain. Specifically, the large time requirement per data set in a TRX experiment is a current limitation for achieving high temporal resolution. Improvements in the speed of data collection will enable a more detailed exploration of the time domain, as reported for T-jump X-ray solution scattering experiments^23^. Additionally, because T-jump induces structural changes that can be distributed across the entire unit cell, rather than concentrated at a specific site (e.g. a chromophore or ligand-binding site), refinement methods that utilize ESFMs do not perform well because there is a significant phase difference between structure factors for the dark state and for the high-energy states that are populated during T-jump^27^. Simultaneous refinement of extrapolated structure factor magnitudes and phases will improve our ability to accurately model high-energy conformations from TRX data. With future technological developments forthcoming, and T-jump available as a general perturbation method, TRX experiments will become more relevant to a wider audience of structural biologists.

## Supporting information

Guide to Supplemental Information

## Methods

### Lysozyme Crystallization

Lysozyme was crystallized in batch by mixing a 20 mg ml^−1^ solution (lyophilized enzyme dissolved in 0.1 M sodium acetate at pH 3.0) with precipitant (28% (w/v) NaCl, 8% (w/v) PEG6000 and 0.1 M sodium acetate at pH 3.0) in a 1:1 ratio as previously described^53^. For inhibitor-bound structures N,N’-diacetylchitobiose was included at 10 mg ml^−1^ with the Lysozyme solution prior to mixture with precipitant. Both apo and chitobiose-bound crystals were approximately 10 μm. Prior to storage, slurries were centrifuged at 4 °C and 3,000g for 3 min, the supernatant was removed and crystals were resuspended in 10% (w/v) NaCl and 1.0 M sodium acetate (pH 3.0).

### Sample Delivery and Data Collection

We collected data at SACLA^35^ using the Diverse Application Platform for Hard X-ray Diffraction in SACLA (DAPHNIS)^54^ at BL2^55^. Prior to data collection, crystal slurries were centrifuged, the supernatant was removed, and crystals were directly mixed (in a 1:9 ratio) with a viscogen containing crystal stabilization buffer with an additional 18% hydroxyethyl cellulose, as previously described^33^. The mixture was then loaded into the reservoir of a sample injector device^34^. This device amplifies pressure generated by an HPLC pump to extrude the viscous medium carrying crystals through a microfluidic nozzle with a diameter of 75 µm. The linear flow rate of this stream was 9.43 mm/s, with a volumetric flow rate of 2.5 µL/min. Helium gas was streamed as a sheath for the liquid jet, with He flow adjusted as needed to maintain a stable stream of lysozyme microcrystals.

X-ray diffraction measurements were made using 0.6 mJ XFEL pulses 10 fs in duration, with a peak photon energy of 9.5 keV. Diffraction images were collected at 30 Hz using a custom-built 4M pixel detector with multi-port CCD (mpCCD) sensors^36^. For time-resolved datasets, crystals were rapidly heated by excitation of the water O-H stretch with mid-IR laser light (1443 nm, 7 ns pulse duration, 540 uJ pulse energy, ∼50 µm (FWHM) focus diameter, (see **Figure EX4**) at defined time delays prior to X-ray exposure. Although the absorption peak at ∼1443 nm is not the strongest mid-IR absorption peak for liquid water, it was selected as a compromise between large T-jumps and uniform heating of the sample, because greater absorption leads to greater increase of thermal energy but also leads to more dramatic temperature gradients through the sample. The IR pump laser operated at a repetition rate of 10 Hz, so data were collected with two dark diffraction measurements interleaved between each T-jump measurement^30, 31^. An IR camera was used to monitor the alignment of the pump laser with the sample, and the firing of the laser was verified by reflecting a small amount of the laser light to a photodiode. Both the jet stream and the pump laser focus were centered on the position of the X-ray beam, which was focused to a diameter of 1.5 µm, much smaller than both the jet and the pump laser. The direction of the IR laser beam was approximately 45° off-axis from the XFEL beam. Data collection parameters were consistent across timepoints and samples (apo vs chitobiose-bound).

### Processing of Bragg Reflections

Data collection was supported by a real-time data processing pipeline^37^ developed on Cheetah^56^. Images identified as hits were processed using methods from CCTBX^38^. For Bragg processing, data were indexed and integrated using *dials.stills_process* ^39^. Initial indexing results were used to refine the detector model, as well as crystal models^40^. Refinement of the detector distance and panel geometry improved the agreement between measured and predicted spots. Bragg data were then merged and post-refined using *cxi.merge* ^41^. Error estimates were treated according to the Ev11 method^40, 57^, wherein error estimates were increased using terms refined from the measurements until they could better explain the uncertainty observed in the merged reflection intensities. A total of 17 merged data sets were produced from our experiment. For the apo enzyme, we collected one “laser off” data set, and explored three pump-probe time delays (20 ns, 20 μs, 200 μs), including a “laser on” data set and two interleaved control data sets (dark1, dark2) for each time delay. For the chitobiose-bound enzyme, we collected one “laser off” data set, and explored two pump-probe time delays (20 ns, 200 μs), with corresponding interleaved control data sets (dark1, dark2).

### Processing and Analysis of Solvent Scattering

For analysis of solvent scattering, each image identified as a hit by Cheetah was processed into an azimuthal averages using *dxtbx.radial_average*. The function was modified to include a sliding one-dimensional median filter, with a window size of 15 q bins. The median filter minimized contributions of the Bragg peaks to the radial average while preserving the solvent scattering signal. Individual scattering curves were culled from later SVD analysis based on collection anomalies (clogging of sample jet, etc.), since these contributions dominated variations in the background signal, making detection of T-jumps difficult. A series of custom scripts were written to process and analyze the solvent scattering curves (https://github.com/fraser-lab/xray_thermometer). Briefly, one-dimensional X-ray scattering curves were pooled and scaled using an algebraic (least-squares) procedure^23^. A mask was applied so that only data between 2.0 Å^-1^and 2.5 Å^-1^ were used for scaling. Scaled curves, containing data between 1.0 Å^-1^ and 2.7 Å^-1^, were combined in a 2D matrix, where each row of the matrix represents a unique scattering curve, each column represents a value of q, and the entries represent the corresponding azimuthally-averaged scattering intensities. This matrix was analyzed using SVD^24, 58–60^, where the right singular vectors (v_n_) represented basis vectors of the observed scattering curves and the left singular vectors (u_n_) described the contribution of each basis vector to each individual scattering curve. By analyzing the values of u_1n_, the entries in U which quantify the contribution of the temperature-dependent singular vector (v_1_) to each scattering curve (n), we determined that u_1n_ correlated with IR laser status.

### Estimation of Temperature Jump

The following calculations assume pure water, which is only an approximation; however, these calculations roughly reproduce the T-jumps that were experimentally measured in prior work^23^. Additionally, we note that the heat capacity of crystalline lysozyme is nearly the same as for pure water^61^.

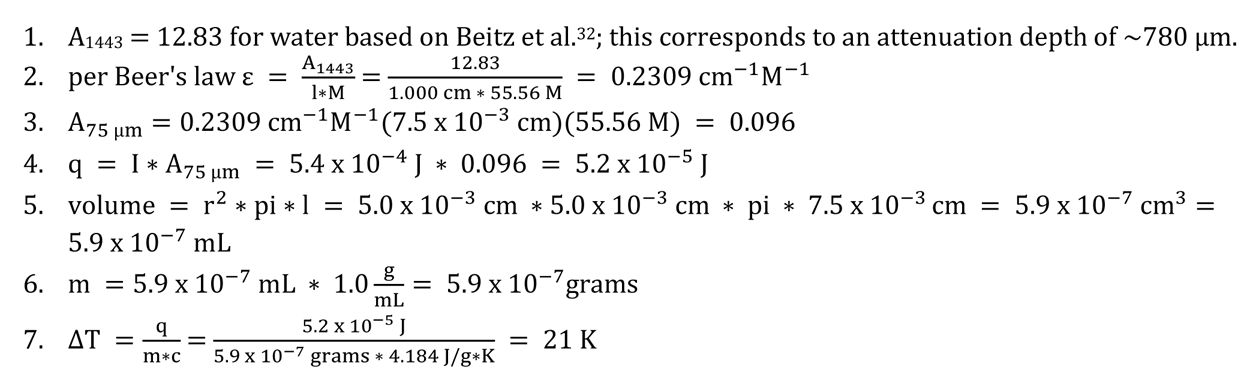

We note that the calculated value of 21 °C likely represents the maximum temperature-jump achieved.

### Initial Model Building

All data as output from *cxi.merge* were converted from intensities to structure factors using *phenix.reflection_file_converter*. Initial phases for each of the 17 data sets (see above) were calculated via MR using Phaser^62^, with PDB entry 1IEE as the search model. R-free flags were carried over from PDB entry 1IEE, and random atomic displacements (0.5 Å) were applied to the coordinates to remove model bias before an initial round of refinement. Iterative model building using Coot followed by refinement in PHENIX^63^ was conducted until the structures converged, following a previously described protocol^64^. The structures refined against the raw F_dark_ and F_t_ structure factors were deposited to the Protein Data Bank). Data collection and refinement statistics for these structures are provided in **Tables EX2-EX5**.

### Calculation of Weighted Difference Electron Density Maps

Data were collected in an interleaved fashion (light, dark1, dark2) as described above, thus for each pump-probe time delay we calculated an (F_t_ – F_dark2_) and control (F_dark1_ – F_dark2_) difference structure factors. Difference structure factors were weighted as previously described^5, 27, 65^ using Reciprocalspaceship^66^.

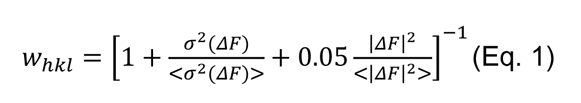

Map phases were calculated using models refined against F_t_.

### Calculation of Integrated Absolute Difference Density Above Threshold (IADDAT)

Finalized difference electron density maps were integrated for the experimental and control datasets for each pump-probe time delay. Briefly, maps were calculated on a common grid using Reciprocalspaceship^66^, with the unit cell sampled using a grid resolution factor corresponding to 0.25*d_min_. Map voxels containing |ρ| ≥ 0.04 e^-^/Å and within 2.5 Å of the atomic model were summed and an average value was calculated per residue. These values were mapped back onto the corresponding structure or plotted directly as a function of residue number for visualization.

### Simulated Difference Density Calculations

To understand how uniform amplification of B-factors might affect difference maps, we simulated data based on the refined laser off structure. The ADPs present in the model were scaled up linearly 20% using *phenix.pdbtools* and stored in a second model file. Structure factors were then calculated for both models using *phenix.fmodel*, and difference maps (F_1.2B_ – F_B_) were created using the custom scripts described above.

### Difference Electron Density Map Correlations

Finalized difference structure factors were created using custom scripts as described above. Structure factors were converted to difference maps using Reciprocalspaceship^66^, maps were sampled on a uniform grid corresponding to 0.25*d_min_ for the first map. Maps were then thresholded at |ρ| ≥ 0.04 e^-^/Å, but map voxels were not “flattened” as a function of distance from the model. Custom Python scripts using Numpy were used to calculate and visualize pairwise correlation coefficients for light and control datasets.

### Modeling High-Energy Conformations and Refinement Against Extrapolated Structure Factor Magnitudes

Extrapolated structure factor magnitudes (ESFMs) were calculated using Reciprocalspaceship^66^ by adding the weighted structure factor differences (see Eq. 1 for the definition of w) back to the dark state structure factors, as previously described^5, 6^.

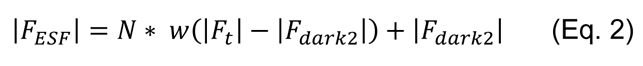

R-free flags were carried over from 1IEE as for the original reflection files. Initial models were prepared for each pump-probe time delay by rigid-body refinement of the laser off structure against ESFMs calculated with N=1. N is the reciprocal of the fraction of molecules occupying the excited conformation, as previously described^27, 28^. Thus, N=1 is not yet extrapolated but represents the experimental T-jumped state with noise down-weighted (see Eq 2. and Figure 1) on a per-structure factor basis (see Eq. 1).

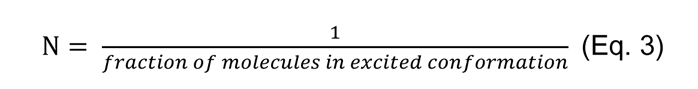

With these initial models of the experimental T-jumped state in hand, alternative conformations representing high-energy states that could be manually interpreted from the difference electron density (e.g. for residues 23 and 97-100, described in the main text) were added to the refined structures. New conformations were labeled as “B” and set to 10% occupancy, while the original refined conformation was set as “A” at 90% occupancy. The resulting multi-conformer structures were then subjected to refinement of coordinates, B-factors, and occupancies against ESFMs calculated with extrapolation factors (N) ranging from 1 to 20. Occupancies of relevant conformations were analyzed as a function of extrapolation factor (N = 1, 2, …,19, 20) using custom Python scripts built upon the GEMMI library (https://github.com/project-gemmi/gemmi). The resulting R-factors from refinement against ESFMs are relatively poor (**Figure EX5**), resulting from inadequacies with the scalar approximation implemented in this procedure^5, 27^, however the stability of the high-energy conformations during the refinements support their accuracy. Due to the generally poor R-factors for these multi-conformer models, we have not deposited them to the PDB; however, they are available on our GitHub page and archived on Zenodo (see below).

### Normal Mode Analysis – Anisotropic Network Model Development

An anisotropic network model (ANM) was developed using a combination of ProDy^67^ and custom Python scripts. Briefly, the model for the 200 µs timepoint was stripped to conformation A (corresponding to the laser off state). The PDB was then imported into Prody, where alpha carbons were selected for residues 1-129. An ANM model was calculated using standard settings (cutoff=15.0 and gamma=1.0). The full ANM model (standard setting of top 20 modes) was then combined into a single mode, where each individual mode was scaled by the associated eigenvalue (λ)

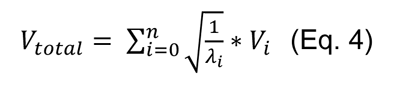

The initial PDB coordinates were then projected along the combined ANM vector and written out to a new PDB file. Finally, the initial and projected coordinates were visualized using PyMOL, with the ANM vectors rendered as arrows between alpha-carbons using the custom module, Modevectors (https://pymolwiki.org/index.php/Modevectors).

### Data Availability

Crystallographic models (refined against the raw structure factors as described above) and the associated reflection data have been deposited in the Protein Data Bank (PDB) under the following accession codes: 8CVU, 8CVV, 8CVW, 8CW0, 8CW1, 8CW3, 8CW5, 8CW6, 8CW7, 8CW8, 8CWB, 8CWC, 8CWD, 8CWE, 8CWF, 8CWG, 8CWH. Additionally, the raw data are publicly available at CXIDB under ID205. All code sufficient to reproduce the analyses described herein, as well as Difference density maps and models refined against ESFMs, are available in a GitHub repository (https://github.com/mthompson-lab/Temperature-Jump-Crystallography_Analysis-for-Paper) and archived on Zenodo (DOI: 10.5281/zenodo.7860140).

## Acknowledgments

The authors thank Takanori Nakane for assistance with real-time data analysis, Sébastien Boutet for advice on experimental design, and Philip Anfinrud for helpful discussions about T-jump experiments. We acknowledge members of the Engineering Team of RIKEN SPring-8 Center for technical support. This work was supported by: grants to MCT and JSF from the NSF BioXFEL Science and Technology Center (STC-1231306); MEXT/JSPS KAKENHI Grants 19H05781 to EN, 19H05784 to MK, and 19H05776 to SI; the Platform Project for Supporting Drug Discovery and Life Science Research (Basis for Supporting Innovative Drug Discovery and Life Science Research) from Japan Agency for Medical Research and Development under Grant JP21am0101070 to SI; and the National Institutes of Health, grant GM117126 to NKS. The XFEL experiments were performed at BL2 of SACLA with the approval of the Japan Synchrotron Radiation Research Institute (JASRI) (Proposal No. 2017B8055 and 2018A8023).

## Author Contributions

MCT and JSF conceptualized the experiments. EN, MK, KT, SI, JMH, and NKS contributed resources and methodology. AMW, EN, IDY, MK, TN, MS, SO, KI, SC, TH, DI, TT, RT, RGS, FY, and MCT conducted investigations. AMW, IDY, ASB, BAB, AB, LJO, NKS, and MCT performed formal analysis of the data. AMW curated the data. JSF, MCT, NKS, EN, MK, and SI administered the project, acquired funding, and supervised research. AMW, MCT, and JSF wrote the manuscript. MCT, AMW, JSF, EN, IDY, BAB, and SC edited the manuscript.

## Correspondence

Correspondence and requests for materials should be addressed to Michael Thompson and Eriko Nango.

## Extended Data (Figures and Tables)

**Figure EX1.**
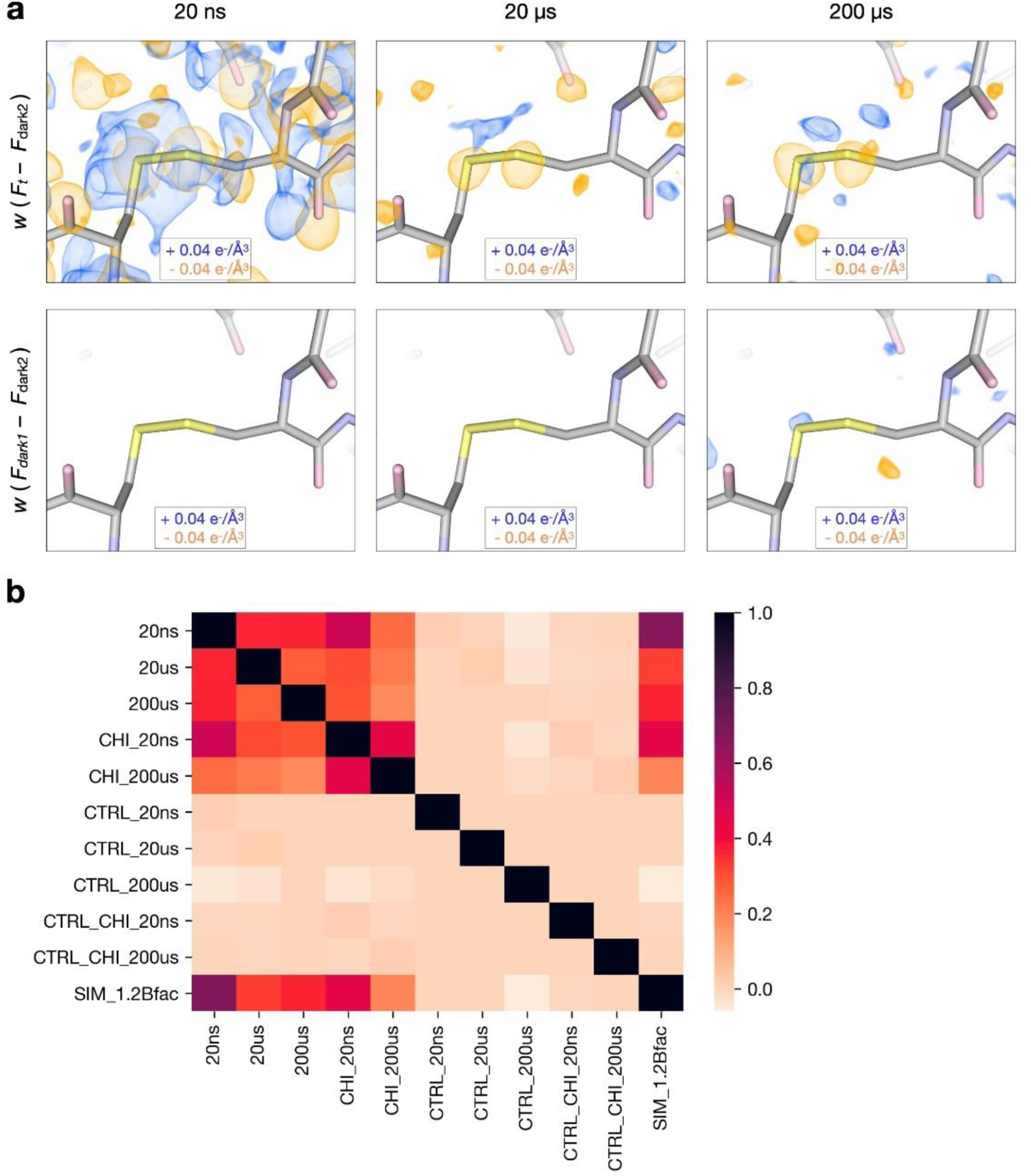
Qualitative and quantitative assessment of time-resolved difference electron density features. (**a**) Comparison of weighted difference density maps for each pump-probe time delay (Flight – Fdark2) and matched controls (Fdark1 – Fdark2) visualized at an absolute contour level of ± 0.04 e^-^/Å^3^ alongside initial refined models. Atoms with greater electron density, such as the disulfide bridge between residues 76 and 94, display clear signals across all experimental maps yet very little noise in matching controls. (**b**) Pairwise correlation coefficients were calculated between all difference maps, revealing varying levels of similarity between experimental maps and low noise across controls. Labels correspond to time-delay (20ns, 20µs, 200µs) presence of the inhibitor, chitobiose (CHI), whether a map was a matched control (CTRL), or based on simulated (SIM) structure factors (see Methods for details).

**Figure EX2.**
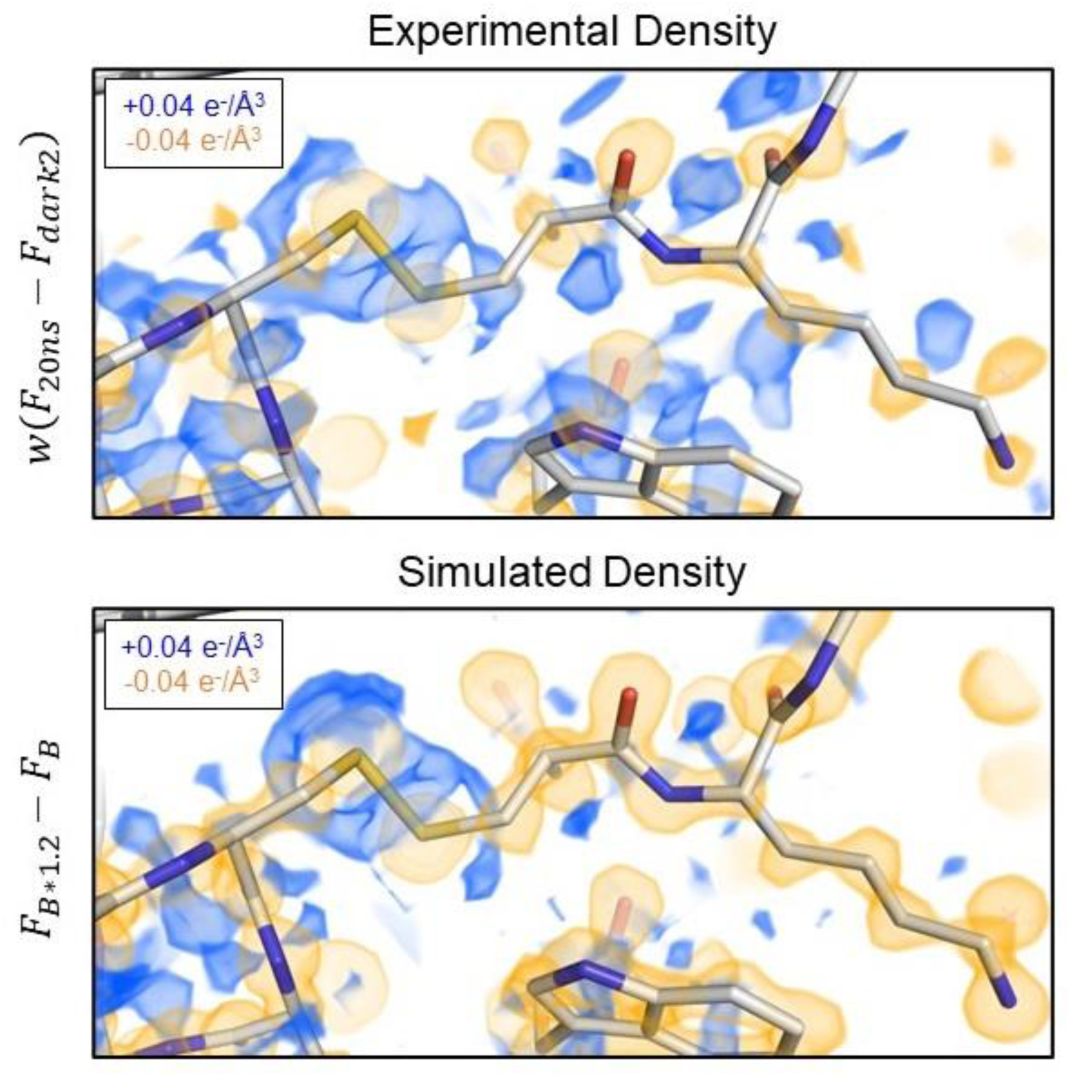
Simulations of increased B-factors recapitulate signals present at the 20 ns pump-probe time delay. The experimental 20ns difference electron density map is visualized along with a simulated difference density map created by linearly scaling the B-factors in the laser off structure by a factor of 1.2. Negative peaks (yellow) are centered upon atoms in both maps, surrounded by positive features (blue).

**Figure EX3.**
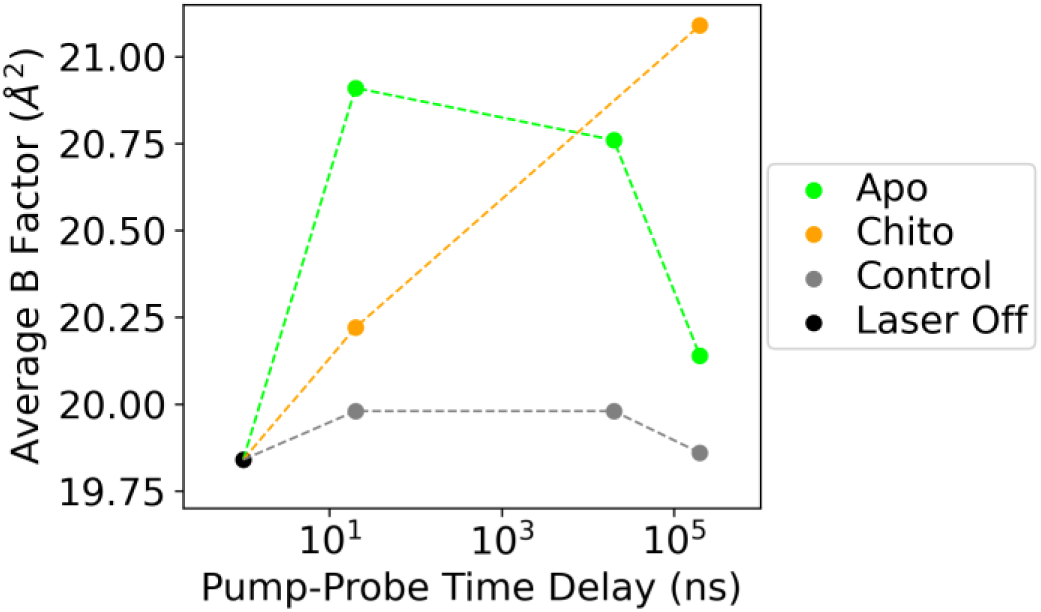
Effect of T-jump on average B-factor of refined apo and chitobiose-bound structures. Models were refined against Laser Off, Experiment (apo or chitobiose bound), or Control structure factors. Controls exhibit similar B-factors across all time points, while B-factors for experimental measurements increase following T-jump. Apo models reveal a decline in B-factors at longer pump-probe time delays as complex motions develop, while chitobiose-bound experimental models retain higher B-factors at 200 μs, indicative of persistent, short-amplitude motions.

**Figure EX4.**
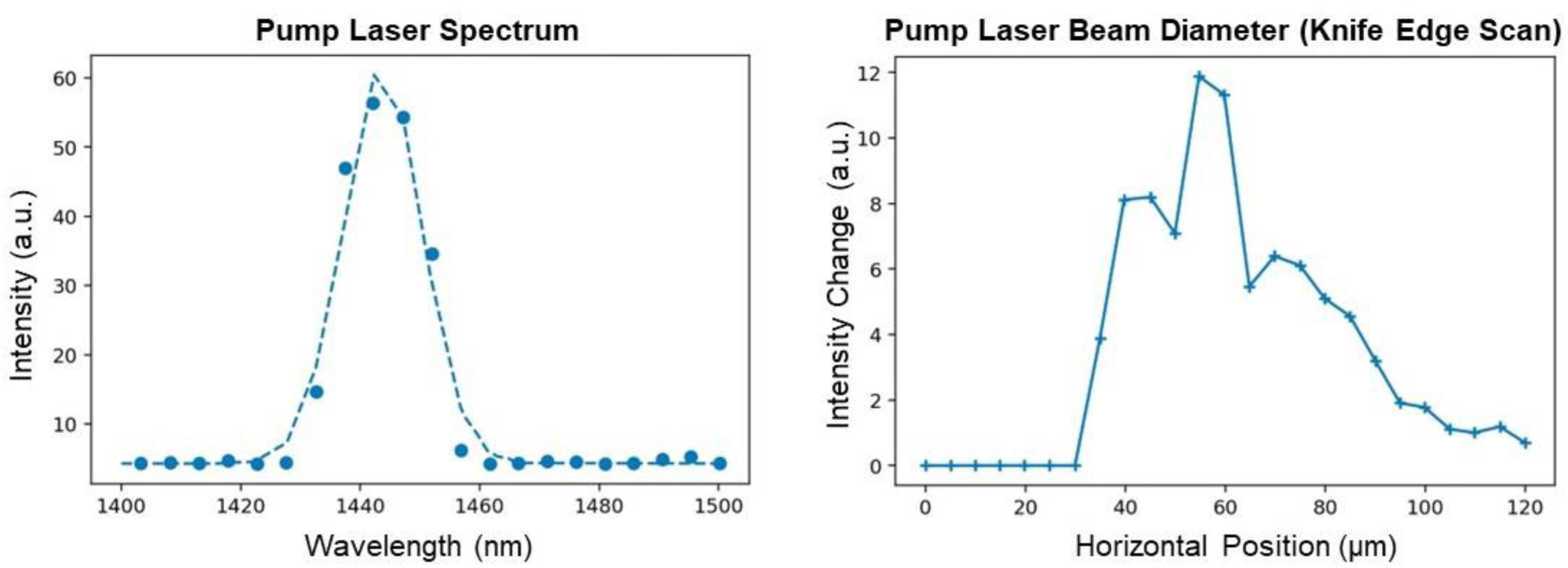
Pump laser parameters. Measurements of pump laser spectrum and beam size (horizontal knife edge scan). The spectrum has a peak at approximately 1443 nm with a bandwidth (FWHM) of approximately 16 nm. The results of a knife edge scan reveal the beam to have an approximate diameter (FWHM) of 50 μm. We note these values represent our best estimations, due to the non-gaussian shape of the plotted data. Nevertheless, the IR beam diameter is much larger than that of the X-ray beam, which is approximately 1.5 μm.

**Figure EX5.**
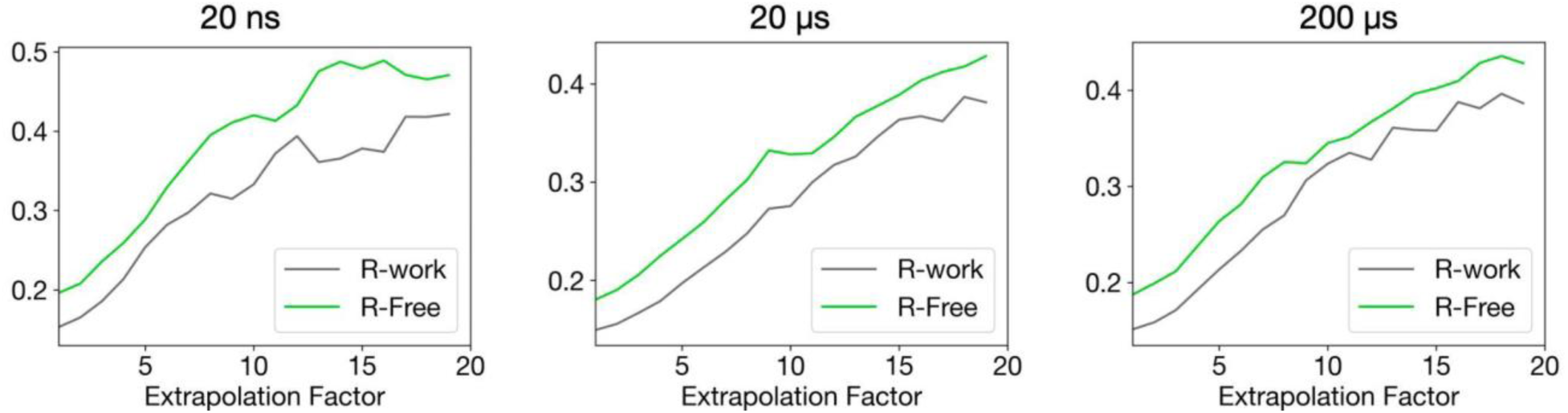
R-factors from extrapolated structure factor refinement. Final R-factors following extrapolated structure factor refinement were plotted as a function of extrapolation factor (N as described in Eq. 2) for the three pump-probe time delays for apo lysozyme structures. Both R-work and R-free increased with extrapolation factor for all three datasets.

**Figure EX6.**
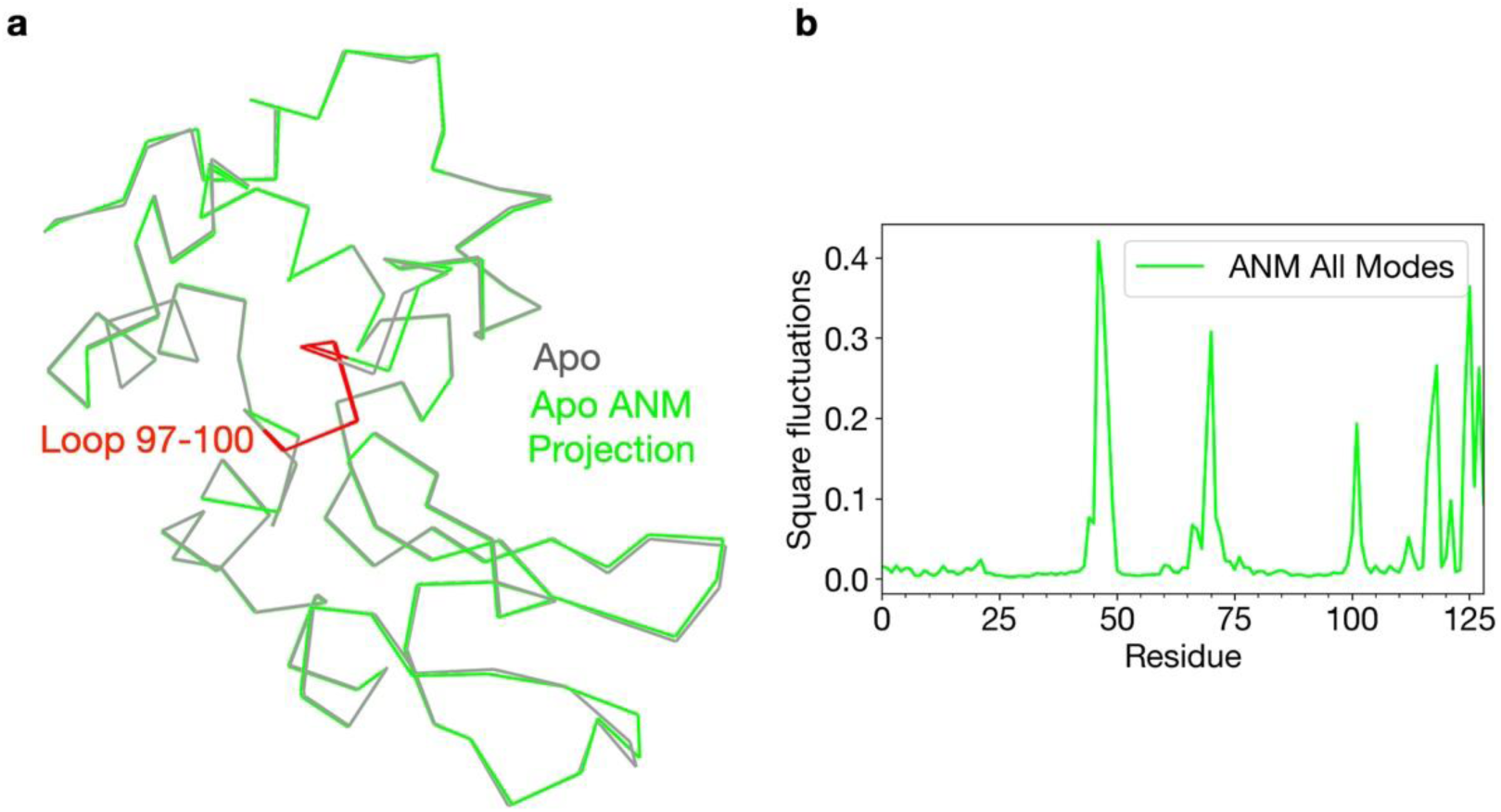
Normal mode analysis of the Apo laser off structure. ProDy was used to generate an anisotropic network model based on the apo ground state conformation. (**a**) The apo structure was then visualized as a ribbon diagram (grey) along with the same model projected along the combined ANM modes (green). (**b**) Per-residue RMSF values for the ANM model were plotted to quantify local dynamics.

**Table EX1.** Sample delivery and X-ray diffraction parameters for apo and chitobiose-bound data collection.

|  |  |
| --- | --- |
| Delivery Matrix | 18% hydroxyethyl cellulose |
| Sample flow rate (μL/min) | 2.5 |
| Capillary diameter (μm) | 75 |
| Linear jet velocity (mm/s) | 9.43 |
| Pressure (psi) | ~3200 |
| X-ray Source | SACLA - BL2 |
| Photon Energy (keV) | 10 |
| X-ray pulse duration (fs) | ~10 |
| Photons per pulse | ~1*10<sup>11</sup> |
| X-ray Repetition Rate (Hz) | 30 |
| IR Pump Laser Wavelength (nm) | 1443 |
| IR Pump Laser Energy (μJ) | 540 |
| IR Pump Laser Pulse Duration (ns) | 7 |

**Table EX2.** Crystallographic statistics for apo data collection. Statistics for the highest-resolution shell are shown in parentheses.

| Dataset | Apo laser off | Light |  |  | Dark1 |  |  | Dark2 |  |  |
| --- | --- | --- | --- | --- | --- | --- | --- | --- | --- | --- |
|  |  | Apo 20ns | Apo 20 us | Apo 200 us | Apo 20ns | Apo 20 us | Apo 200 us | Apo 20ns | Apo 20 us | Apo 200 us |
| Hit rate (%) | 22.1 | 35.2 | 39.2 | 27.0 | 35.9 | 39.8 | 27.1 | 36.3 | 40.5 | 27.4 |
| Indexing rate (%) | 84.7 | 91.7 | 87.8 | 82.8 | 87.9 | 87.0 | 81.6 | 88.1 | 87.2 | 81.3 |
| No. of images | 9464 | 15253 | 13681 | 11931 | 14383 | 13559 | 11857 | 14454 | 13893 | 11950 |
| Resolution Range | 28.09-1.57<br>(1.60-1.57) | 31.62 -1.57<br>(1.60-1.57) | 28.11-1.57<br>(1.60-1.57) | 28.11-1.57<br>(1.60-1.57) | 28.09 -1.57<br>(1.60-1.57) | 28.09 -1.57<br>(1.60-1.57) | 28.09 -1.65<br>(1.68-1.65) | 28.09 -1.57<br>(1.60-1.57) | 28.09 -1.57<br>(1.60-1.57) | 28.09 -1.57<br>(1.60-1.57) |
| Space Group | P4<sub>3</sub>2<sub>1</sub>2 | P4<sub>3</sub>2<sub>1</sub>2 | P4<sub>3</sub>2<sub>1</sub>2 | P4<sub>3</sub>2<sub>1</sub>2 | P4<sub>3</sub>2<sub>1</sub>2 | P4<sub>3</sub>2<sub>1</sub>2 | P4<sub>3</sub>2<sub>1</sub>2 | P4<sub>3</sub>2<sub>1</sub>2 | P4<sub>3</sub>2<sub>1</sub>2 | P4<sub>3</sub>2<sub>1</sub>2 |
| Unit-cell parameters |  |  |  |  |  |  |  |  |  |  |
| a (Å) | 79.45 +/- 0.10 | 79.52 +/- 0.13 | 79.52 +/- 0.10 | 79.51 +/- 0.11 | 79.44 +/- 0.10 | 79.44 +/- 0.09 | 79.45 +/- 0.10 | 79.44 +/- 0.09 | 79.44 +/- 0.09 | 79.46 +/- 0.11 |
| b (Å) | 79.45 +/- 0.10 | 79.52 +/- 0.13 | 79.52 +/- 0.10 | 79.51 +/- 0.11 | 79.44 +/- 0.10 | 79.44 +/- 0.09 | 79.45 +/- 0.10 | 79.44 +/- 0.09 | 79.44 +/- 0.09 | 79.46 +/- 0.11 |
| c (Å) | 38.16 +/- 0.07 | 38.23 +/- 0.08 | 38.21 +/- 0.05 | 38.19 +/- 0.05 | 38.19 +/- 0.06 | 38.20 +/- 0.05 | 38.19 +/- 0.06 | 38.19 +/- 0.06 | 38.20 +/- 0.06 | 38.19 +/- 0.06 |
| alpha (°) | 90 | 90 | 90 | 90 | 90 | 90 | 90 | 90 | 90 | 90 |
| beta (°) | 90 | 90 | 90 | 90 | 90 | 90 | 90 | 90 | 90 | 90 |
| gamma (°) | 90 | 90 | 90 | 90 | 90 | 90 | 90 | 90 | 90 | 90 |
| Total Reflections | 1593940<br>(39912) | 3195992<br>(85262) | 2208287<br>(56144) | 1783773<br>(45477) | 2279135<br>(58015) | 2144941<br>(54395) | 1601963<br>(48482) | 2302697<br>(58593) | 2202111<br>(55792) | 1732358<br>(43260) |
| Multiplicity | 90.55 (46.63) | 181.56<br>(99.61) | 125.46<br>(65.59) | 101.34<br>(53.13) | 129.48<br>(67.77) | 121.86<br>(63.55) | 105.36<br>(67.43) | 130.82 (68.45) | 125.11 (65.18) | 98.41 (50.54) |
| Completeness (%) | 99.97 (100) | 99.98<br>(100.00) | 99.97<br>(100.00) | 99.97<br>(100.00) | 99.97 (100) | 99.97 (100) | 99.97 (100) | 99.97 (100) | 99.97 (100) | 99.98 (100) |
| Mean I/sigma (I) | 9.463 (5.889) | 12.969<br>(6.918) | 10.844<br>(6.633) | 9.986 (6.280) | 10.619<br>(6.240) | 10.000<br>(6.040) | 9.772 (6.896) | 10.612 (6.120) | 10.580 (6.353) | 9.384 (6.167) |
| Wilson B (Å<sup>2</sup>) | 14.1 | 15.37 | 15.38 | 15.1 | 15.23 | 15.30 | 14.807 | 15.26 | 15.38 | 14.84 |
| R-split (%) | 9.4 (40.0) | 5.9 (25.9) | 7.5 (25.7) | 8.5 (30.6) | 7.8 (30.5) | 8.4 (32.1) | 9.9 (16.6) | 8.1 (25.9) | 7.9 (30.4) | 9.2 (28.5) |
| CC-int (%) | 98.6 (17.0) | 99.6 (34.6) | 99.2 (44.5) | 98.4 (37.7) | 99.1 (39.7) | 99.0 (31.9) | 71.7 (81.9) | 98.6 (56.0) | 99.1 (55.0) | 96.9 (44.3) |

**Table EX3.**
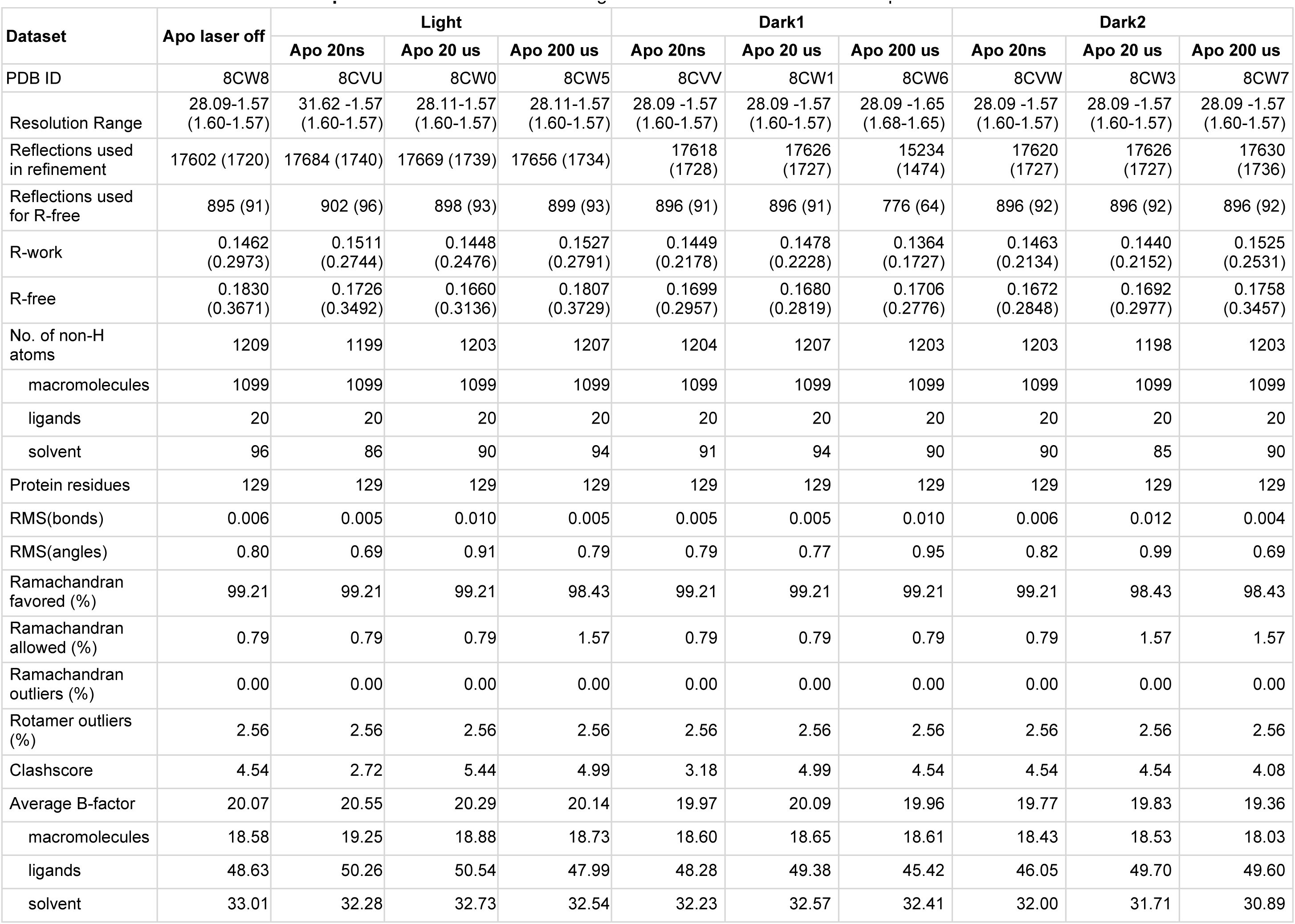
Refinement statistics for apo datasets. Statistics for the highest-resolution shell are shown in parentheses.

| Dataset | Apo laser off | Light |  |  | Dark1 |  |  | Dark2 |  |  |
| --- | --- | --- | --- | --- | --- | --- | --- | --- | --- | --- |
|  |  | Apo 20ns | Apo 20 us | Apo 200 us | Apo 20ns | Apo 20 us | Apo 200 us | Apo 20ns | Apo 20 us | Apo 200 us |
| PDB ID | 8CW8 | 8CVU | 8CW0 | 8CW5 | 8CVV | 8CW1 | 8CW6 | 8CVW | 8CW3 | 8CW7 |
| Resolution Range | 28.09-1.57<br>(1.60-1.57) | 31.62 -1.57<br>(1.60-1.57) | 28.11-1.57<br>(1.60-1.57) | 28.11-1.57<br>(1.60-1.57) | 28.09 -1.57<br>(1.60-1.57) | 28.09 -1.57<br>(1.60-1.57) | 28.09 -1.65<br>(1.68-1.65) | 28.09 -1.57<br>(1.60-1.57) | 28.09 -1.57<br>(1.60-1.57) | 28.09 -1.57<br>(1.60-1.57) |
| Reflections used in refinement | 17602 (1720) | 17684 (1740) | 17669 (1739) | 17656 (1734) | 17618 (1728) | 17626 (1727) | 15234 (1474) | 17620 (1727) | 17626 (1727) | 17630 (1736) |
| Reflections used for R-free | 895 (91) | 902 (96) | 898 (93) | 899 (93) | 896 (91) | 896 (91) | 776 (64) | 896 (92) | 896 (92) | 896 (92) |
| R-work | 0.1462<br>(0.2973) | 0.1511<br>(0.2744) | 0.1448<br>(0.2476) | 0.1527<br>(0.2791) | 0.1449<br>(0.2178) | 0.1478<br>(0.2228) | 0.1364<br>(0.1727) | 0.1463<br>(0.2134) | 0.1440<br>(0.2152) | 0.1525<br>(0.2531) |
| R-free | 0.1830<br>(0.3671) | 0.1726<br>(0.3492) | 0.1660<br>(0.3136) | 0.1807<br>(0.3729) | 0.1699<br>(0.2957) | 0.1680<br>(0.2819) | 0.1706<br>(0.2776) | 0.1672<br>(0.2848) | 0.1692<br>(0.2977) | 0.1758<br>(0.3457) |
| No. of non-H atoms | 1209 | 1199 | 1203 | 1207 | 1204 | 1207 | 1203 | 1203 | 1198 | 1203 |
| macromolecules | 1099 | 1099 | 1099 | 1099 | 1099 | 1099 | 1099 | 1099 | 1099 | 1099 |
| ligands | 20 | 20 | 20 | 20 | 20 | 20 | 20 | 20 | 20 | 20 |
| solvent | 96 | 86 | 90 | 94 | 91 | 94 | 90 | 90 | 85 | 90 |
| Protein residues | 129 | 129 | 129 | 129 | 129 | 129 | 129 | 129 | 129 | 129 |
| RMS(bonds) | 0.006 | 0.005 | 0.010 | 0.005 | 0.005 | 0.005 | 0.010 | 0.006 | 0.012 | 0.004 |
| RMS(angles) | 0.80 | 0.69 | 0.91 | 0.79 | 0.79 | 0.77 | 0.95 | 0.82 | 0.99 | 0.69 |
| Ramachandran favored (%) | 99.21 | 99.21 | 99.21 | 98.43 | 99.21 | 99.21 | 99.21 | 99.21 | 98.43 | 98.43 |
| Ramachandran allowed (%) | 0.79 | 0.79 | 0.79 | 1.57 | 0.79 | 0.79 | 0.79 | 0.79 | 1.57 | 1.57 |
| Ramachandran outliers (%) | 0.00 | 0.00 | 0.00 | 0.00 | 0.00 | 0.00 | 0.00 | 0.00 | 0.00 | 0.00 |
| Rotamer outliers (%) | 2.56 | 2.56 | 2.56 | 2.56 | 2.56 | 2.56 | 2.56 | 2.56 | 2.56 | 2.56 |
| Clashscore | 4.54 | 2.72 | 5.44 | 4.99 | 3.18 | 4.99 | 4.54 | 4.54 | 4.54 | 4.08 |
| Average B-factor | 20.07 | 20.55 | 20.29 | 20.14 | 19.97 | 20.09 | 19.96 | 19.77 | 19.83 | 19.36 |
| macromolecules | 18.58 | 19.25 | 18.88 | 18.73 | 18.60 | 18.65 | 18.61 | 18.43 | 18.53 | 18.03 |
| ligands | 48.63 | 50.26 | 50.54 | 47.99 | 48.28 | 49.38 | 45.42 | 46.05 | 49.70 | 49.60 |
| solvent | 33.01 | 32.28 | 32.73 | 32.54 | 32.23 | 32.57 | 32.41 | 32.00 | 31.71 | 30.89 |

**Table EX4.** Crystallographic statistics for chitobiose-bound data collection. Statistics for the highest-resolution shell are shown in parentheses.

| Dataset | Chito laser off | Light |  | Dark1 |  | Dark2 |  |
| --- | --- | --- | --- | --- | --- | --- | --- |
|  |  | Chito 20ns | Chito 200us | Chito 20ns | Chito 200us | Chito 20ns | Chito 200us |
| Hit rate (%) | 25.3 | 30.2 | 32.3 | 32.0 | 32.7 | 32.2 | 32.9 |
| Indexing rate (%) | 89.3 | 92.4 | 93.7 | 91.1 | 92.9 | 90.7 | 93.0 |
| No. of images | 31575 | 22549 | 21472 | 22489 | 20882 | 21715 | 21067 |
| Resolution Range | 30.74 -1.51<br>(1.54-1.51) | 30.74 -1.48<br>(1.51-1.48) | 30.74-1.50<br>(1.53-1.50) | 30.74 -1.48<br>(1.51-1.48) | 30.74-1.50<br>(1.53-1.50) | 30.74 -1.48<br>(1.51-1.48) | 30.74-1.50<br>(1.53-1.50) |
| Space Group | P4<sub>3</sub>2<sub>1</sub>2 | P4<sub>3</sub>2<sub>1</sub>2 | P4<sub>3</sub>2<sub>1</sub>2 | P4<sub>3</sub>2<sub>1</sub>2 | P4<sub>3</sub>2<sub>1</sub>2 | P4<sub>3</sub>2<sub>1</sub>2 | P4<sub>3</sub>2<sub>1</sub>2 |
| Unit-cell parameters |  |  |  |  |  |  |  |
| a (Å) | 78.55 +/- 0.22 | 79.05 +/- 0.16 | 78.98 +/- 0.12 | 78.91 +/- 0.11 | 78.85 +/- 0.09 | 78.91 +/- 0.11 | 78.85 +/- 0.09 |
| b (Å) | 78.55 +/- 0.22 | 79.05 +/- 0.16 | 78.98 +/- 0.12 | 78.91 +/- 0.11 | 78.85 +/- 0.09 | 78.91 +/- 0.11 | 78.85 +/- 0.09 |
| c (Å) | 38.30 +/- 0.13 | 38.50 +/- 0.05 | 38.47 +/- 0.04 | 38.48 +/- 0.05 | 38.51 +/- 0.04 | 38.48 +/- 0.05 | 38.51 +/- 0.04 |
| alpha (°) | 90 | 90 | 90 | 90 | 90 | 90 | 90 |
| beta (°) | 90 | 90 | 90 | 90 | 90 | 90 | 90 |
| gamma (°) | 90 | 90 | 90 | 90 | 90 | 90 | 90 |
| Total Reflections | 10094940 (25562) | 3815383 (30777) | 3053934 (31769) | 3283906 (33942) | 2892458 (34076) | 3127348 (31601) | 2895230 (33351) |
| Multiplicity | 556.53 (28.92) | 198.25 (32.95) | 165.09 (35.54) | 170.64 (36.34) | 156.36 (38.12) | 162.50 (33.83) | 156.51 (37.31) |
| Completeness (%) | 99.98 (100) | 99.98 (100) | 99.98 (100) | 99.98 (100) | 99.98 (100) | 99.98 (100) | 99.98 (100) |
| Mean I/sigma (I) | 18.535 (3.367) | 12.251 (3.347) | 11.487 (3.673) | 11.746 (3.977) | 11.275 (3.834) | 11.287 (3.831) | 11.342 (3.883) |
| Wilson B (Å<sup>2</sup>) | 14.51 | 16.16 | 16.40 | 15.03 | 15.82 | 14.97 | 15.76 |
| R-split (%) | 3.5 (23.1) | 5.5 (30.7) | 5.7 (25.8) | 6.1 (25.6) | 6.1 (25.4) | 6.2 (25.0) | 6.2 (25.0) |
| CC-int (%) | 99.9 (89.3) | 99.6 (78.7) | 99.5 (86.7) | 99.3 (84.1) | 99.4 (83.5) | 99.4 (82.3) | 99.4 (86.0) |

**Table EX5.** Refinement statistics for chitobiose-bound datasets. Statistics for the highest-resolution shell are shown in parentheses.

| Dataset | Chito laser off | Light |  | Dark1 |  | Dark2 |  |
| --- | --- | --- | --- | --- | --- | --- | --- |
|  |  | Chito 20ns | Chito 200us | Chito 20ns | Chito 200us | Chito 20ns | Chito 200us |
| PDB ID | 8CWB | 8CWC | 8CWF | 8CWD | 8CWG | 8CWE | 8CWH |
| Resolution Range | 30.74 -1.51<br>(1.54-1.51) | 30.74 -1.48<br>(1.51-1.48) | 30.74-1.50<br>(1.53-1.50) | 30.74 -1.48<br>(1.51-1.48) | 30.74-1.50<br>(1.53-1.50) | 30.74 -1.48<br>(1.51-1.48) | 30.74-1.50<br>(1.53-1.50) |
| Reflections used in refinement | 18138 (1754) | 19245 (1873) | 18499 (1825) | 19245 (1873) | 18499 (1825) | 19245 (1873) | 18499 (1825) |
| Reflections used for R-free | 930 (97) | 984 (103) | 950 (99) | 984 (103) | 950 (99) | 984 (103) | 950 (99) |
| R-work | 0.1357 (0.1329) | 0.1316 (0.1246) | 0.1332 (0.1105) | 0.1306 (0.1147) | 0.1292 (0.1066) | 0.1298 (0.1137) | 0.1321 (0.1051) |
| R-free | 0.1679 (0.2134) | 0.1683 (0.2273) | 0.1660 (0.2130) | 0.1633 (0.2039) | 0.1663 (0.2210) | 0.1632 (0.2090) | 0.1679 (0.1996) |
| No. of non-H atoms | 1157 | 1155 | 1153 | 1162 | 1160 | 1161 | 1166 |
| macromolecules | 1030 | 1030 | 1030 | 1030 | 1030 | 1030 | 1030 |
| ligands | 64 | 64 | 64 | 64 | 64 | 64 | 64 |
| solvent | 91 | 89 | 87 | 96 | 94 | 95 | 100 |
| Protein residues | 129 | 129 | 129 | 129 | 129 | 129 | 129 |
| RMS(bonds) | 0.018 | 0.016 | 0.019 | 0.018 | 0.018 | 0.019 | 0.020 |
| RMS(angles) | 1.21 | 1.19 | 1.20 | 1.19 | 1.17 | 1.23 | 1.21 |
| Ramachandran favored (%) | 99.21 | 99.21 | 99.21 | 99.21 | 99.21 | 99.21 | 99.21 |
| Ramachandran allowed (%) | 0.79 | 0.79 | 0.79 | 0.79 | 0.79 | 0.79 | 0.79 |
| Ramachandran outliers (%) | 0.00 | 0.00 | 0.00 | 0.00 | 0.00 | 0.00 | 0.00 |
| Rotamer outliers (%) | 0.00 | 0.00 | 0.92 | 0.92 | 0.92 | 0.00 | 0.92 |
| Clashscore | 3.87 | 5.80 | 5.32 | 3.87 | 5.80 | 3.38 | 5.80 |
| Average B-factor | 19.58 | 21.08 | 20.97 | 19.42 | 20.67 | 19.57 | 20.39 |
| macromolecules | 18.53 | 19.98 | 19.87 | 18.19 | 19.59 | 18.30 | 19.05 |
| ligands | 22.63 | 26.70 | 26.06 | 24.83 | 24.94 | 25.34 | 25.04 |
| solvent | 30.23 | 31.55 | 31.91 | 30.55 | 30.90 | 31.21 | 32.55 |

