## Supplementary material for "Mapping Protein Dynamics at High Spatial Resolution with Temperature-Jump X-ray Crystallography": Guide to Supplemental Information

*Supplemental Video S1*
360° view of difference electron density map shown in figure 3A corresponding to 20 ns pump-probe time delay (left panel).

*Supplemental Video S2*
360° view of difference electron density map shown in figure 3A corresponding to 20 µs pump-probe time delay (center panel).

*Supplemental Video S3*
360° view of difference electron density map shown in figure 3A corresponding to 200 µs pump-probe time delay (center panel).

*Supplemental Video S4*
360° view of experimentally-determined difference electron density map calculated for 20 ns pump-probe time delay.

*Supplemental Video S5*
360° view of simulated difference electron density map calculated between dark state model and a model with elevated B-factors.

*Supplemental Video S6*
360° view of difference electron density around residue Y23, calculated for 20 ns pump-probe time delay and shown in figure 4A (left panel).

*Supplemental Video S7*
360° view of difference electron density around the loop containing residues 97-100, calculated for 200 µs pump-probe time delay and shown in figure 4A (right panel).
